# High-resolution 3D genome mapping of late replicative senescence reveals hypomethylation-associated loop expansion, weakened cell identity, and locus-specific L1 derepression

**DOI:** 10.1101/2025.06.11.659151

**Authors:** Maxfield M.G. Kelsey, Audrey Dalgarno, Shane A. Evans, Thomas A. Nunez, Azucena Rocha, Kelly Clark, John M. Sedivy, Nicola Neretti

**Affiliations:** Center on the Biology of Aging, Brown University, Providence, RI, USA; Department of Molecular Biology, Cell Biology, and Biochemistry, Brown University, Providence, RI, USA; Center for Computational Molecular Biology, Brown University, Providence, RI, USA

## Abstract

Cellular senescence contributes to aging and age-associated diseases through the senescence-associated secretory phenotype (SASP), which is fueled in part by derepression of retrotransposable elements. While senescence is driven by transcriptional and epigenetic changes, the contribution of higher-order genome organization remains incompletely defined, particularly at the resolution required to detect chromatin loops. Here we present ∼3 kb-resolution Hi-C maps of late-stage replicatively senescent (RS), proliferating, and quiescent human lung fibroblasts, integrated with total RNA-seq and Nanopore long-read DNA sequencing. RS cells exhibit widespread compartment and subcompartment switching toward active states, consistent with a global loss of heterochromatin. We observe a striking expansion of chromatin loops in RS cells, with an approximately sixfold excess of senescence-specific loops over proliferation– or quiescence-specific loops. These senescence-specific loops are enriched in the B compartment relative to other loop classes, are accompanied by DNA hypomethylation at loop anchors, and are enriched for neuronal-lineage gene programs that are concordantly upregulated, suggesting a structural correlate of the loss of cell-type identity recently described in aging chromatin. Compartment and subcompartment reorganization, but not loop status, is associated with retrotransposon derepression. We identify L1HS_14q23.2_3, an intact L1HS element, as a locus where compartment switching, subcompartment activation, novel loop formation, and pronounced promoter demethylation appear to promote locus-specific derepression. Quiescent fibroblasts unexpectedly share inflammatory transcriptional features with RS cells, raising important considerations for experimental design in senescence research.

## Introduction

Cellular senescence is a stable form of proliferative arrest, typically triggered by irreparable DNA damage from telomere shortening, oncogenic stress, or irradiation (López-Otín et al. 2013). While senescence plays beneficial roles in embryogenesis (Muñoz-Espín et al. 2013), wound healing (Demaria et al. 2014), and tumor suppression (Coppé et al. 2010), it also drives age-associated dysfunction: senescent cells secrete proinflammatory factors collectively known as the senescence-associated secretory phenotype (SASP), which can promote tumorigenesis and induce secondary senescence in neighboring cells (Acosta et al. 2013; Teo et al. 2019; Coppé et al. 2010), and clearance of senescent cells in mice extends lifespan and healthspan (Baker et al. 2011, 2016). Senescence is also accompanied by widespread epigenetic remodeling at every level of chromatin organization (Rocha et al. 2022), including derepression of retrotransposable elements (RTEs). RTE derepression is now recognized as a direct driver of the SASP: cytosolic accumulation of LINE-1 (L1) cDNA in late senescent cells activates the cGAS–STING DNA-sensing pathway and triggers a type I interferon response that reinforces the inflammatory secretome (De Cecco et al. 2019), and L1 derepression has been shown to drive sterile inflammation in aged tissues in vivo (Simon et al. 2019).

Chromosome conformation capture technologies such as Hi-C have revealed that nuclear DNA is organized hierarchically across multiple scales (Lieberman-Aiden et al. 2009). At the coarsest level, the genome is partitioned into open, transcriptionally active A compartments and closed, repressed B compartments. These are further subdivided into subcompartments with distinct long-range interaction patterns and characteristic epigenetic marks (Rao et al. 2014). As Hi-C resolution has improved, individual chromatin loops have also been resolved; these loops can connect promoters and enhancers and are thought to form through a loop extrusion process involving CCCTC-binding factor (CTCF) and cohesin (Sanborn et al. 2015; Fudenberg et al. 2016).

Senescent cells undergo coordinated changes in nuclear structure, including increased cell size (Zhao and Darzynkiewicz 2013) and breakdown of the nuclear lamina (Sadaie et al. 2013; Shah et al. 2013). Several studies have examined how these nuclear changes manifest in 3D genome organization (Chandra et al. 2015; Criscione, De Cecco, et al. 2016; Iwasaki et al. 2019; Olan et al. 2020; Sati et al. 2020; Zirkel et al. 2018), reporting alterations in long– and short-range contacts as well as switching between the A and B compartments, with consequences for chromosome shape and gene regulation. In oncogene-induced senescence (OIS), de novo cohesin binding has been shown to redistribute chromatin loops (Olan et al. 2020). A first limitation of prior work concerns resolution: senescence Hi-C maps have been limited to approximately 10 kb or coarser (Olan et al. 2020), restricting their ability to resolve loops and the finer features of subcompartment organization.

A second limitation is that the late stage of senescence has been underrepresented in prior work. Senescence is a dynamic process that progresses through distinct stages, from “early” to “late”, as cells undergo further changes including lamin B1 downregulation, continued chromatin remodeling, and progressive retrotransposon activation (van Deursen 2014). The late stage is often underrepresented in cell culture studies because of the technical challenges of maintaining cells in a senescent state over extended periods. Yet during aging, the failure to clear senescent cells in vivo means that most senescent cells encountered in tissue are likely in the late stage, making this the most relevant stage for understanding age-associated pathology.

A third limitation is that the 3D-genomic basis of a recently recognized feature of the senescent state, a partial loss of cell-type identity, remains unexamined. Work in aging models, most notably the ICE (Inducible Changes to the Epigenome) system, has shown that loss of epigenetic information drives ectopic activation of lineage-inappropriate gene programs and a structural blurring of cellular identity (Yang et al. 2023). Analogous identity-loss phenotypes have been reported across aging tissues, including transcriptomic convergence between distinct organs in old animals (Izgi et al. 2022) and a gradual erosion of cell-type-defining expression programs in the aging immune compartment (Connolly et al. 2024). In senescent cells specifically, partial loss of pre-senescence identity has now been documented at single-cell resolution, with senescent cells of multiple lineages co-opting features of unrelated cell types or developmental programs (Hernandez-Segura et al. 2017; Huna et al. 2024). Whether this identity drift has a structural basis in the 3D genome, however, has not been systematically examined. Of particular interest are neuronal-lineage programs: senescent astrocytes and keratinocytes ectopically upregulate “neurogenesis” and “neuron differentiation” gene sets (Hernandez-Segura et al. 2017), senescent human fibroblasts induce the neural adhesion molecule L1CAM at the cell surface (Mrazkova et al. 2018), and fibroblasts have long been known to lie close to neurons in the developmental epigenetic landscape, a proximity exploited by direct-conversion protocols in which only three transcription factors (ASCL1, BRN2, MYT1L) are sufficient to convert fibroblasts to functional neurons (Wapinski et al. 2013; Vierbuchen et al. 2010).

## Results

### High-Resolution Hi-C Mapping of Replicative Senescence

To investigate how 3D genome organization supports cellular senescence, we performed in situ Hi-C on proliferating (PRO) and late-stage replicative senescence (RS) LF1 lung fibroblasts, with three biological replicates per condition. Replicates clustered closely in the multidimensional scaling (MDS) plot, demonstrating high reproducibility and allowing us to combine them into the most deeply sequenced Hi-C maps of senescent cells to date, achieving a resolution of 2.75 kb for RS cells (Figure S1, Figure S2, Supplementary Table 1). Proliferating controls reached a resolution of 3.35 kb.

### Compartment Switching in RS

Previous studies have reported a bias toward compartment switching from the transcriptionally repressed B compartment to the active A compartment during senescence (Iwasaki et al. 2019). To compare our RS data with another senescence paradigm, we re-analyzed the highest-resolution (∼10 kb) publicly available Hi-C dataset of OIS which examined HRAS-G12V expressing IMR-90 lung fibroblasts (Olan et al. 2020). Both RS and OIS exhibited a pronounced asymmetry in compartment switching, with substantially more of the genome transitioning toward A than toward B (Figure 1A). In RS, 5.1% of the genome shifted toward A while only 2.4% shifted toward B; OIS showed a similar but slightly attenuated pattern (4.6% toward A, 2.1% toward B; chi-square test, p < 0.001 for both conditions). These results are consistent with the loss-of-heterochromatin model of aging and senescence (Villeponteau 1997), which posits that global erosion of repressive chromatin promotes transcriptional derepression (De Cecco, Criscione, Peckham, et al. 2013). Because compartment scores are normalized within each sample, A/B assignments capture the redistribution of chromatin between active and inactive states rather than absolute changes in compartment volume. Consequently, a directional asymmetry in switching — as observed here in both RS and OIS, and reported previously (Iwasaki et al. 2019) — is consistent with a loss of heterochromatin but cannot, on its own, distinguish weakening of the B compartment from strengthening of the A compartment.

**Figure 1.**
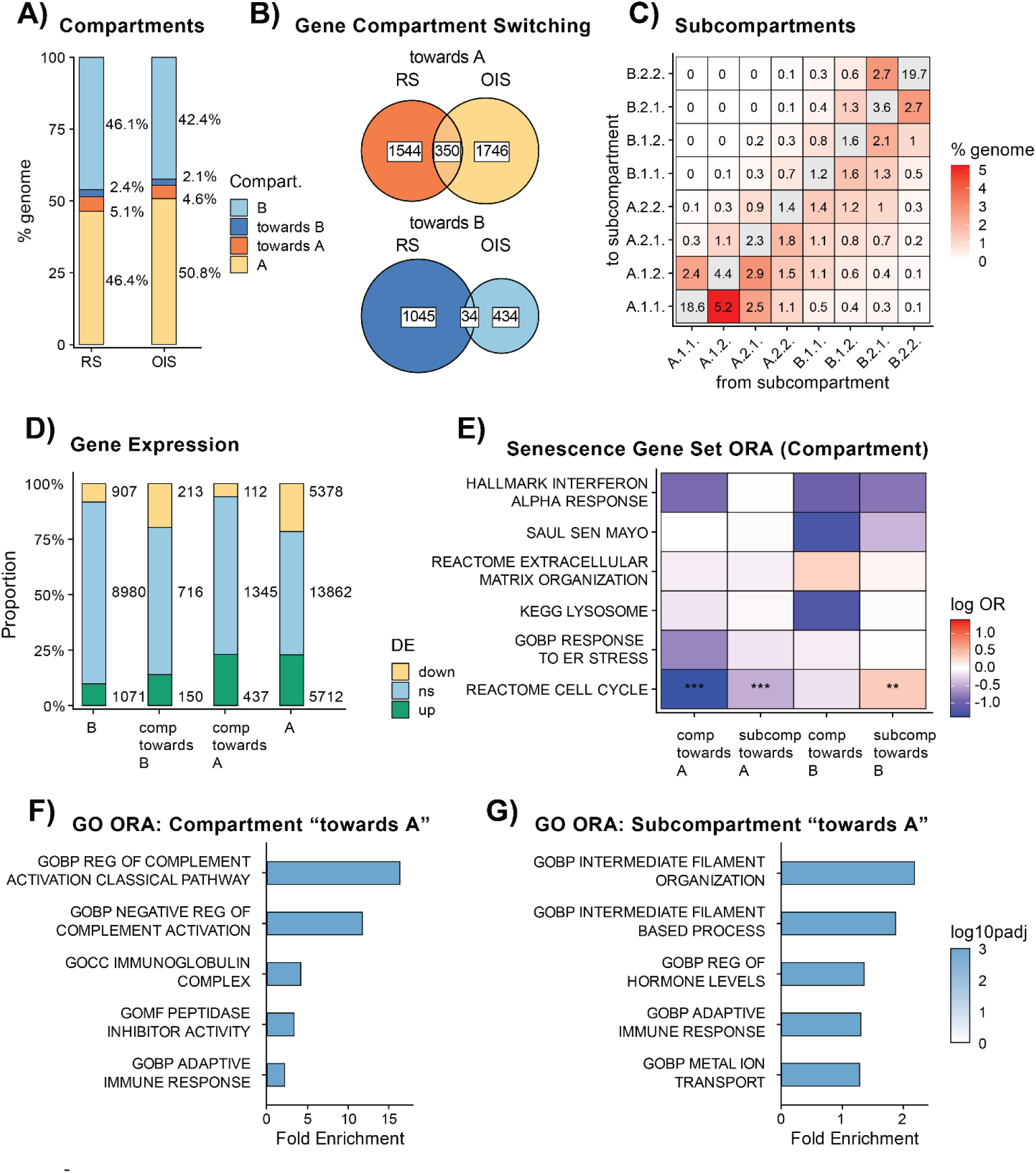
| Compartment and subcompartment switching in RS indicate a moderate global loss of heterochromatin. (**A**) Distribution of the genome across compartment states in replicative senescence (RS) and oncogene-induced senescence (OIS). (**B**) Overlap between genes undergoing compartment switching toward A (top) or toward B (bottom) in RS and OIS. (**C**) Percentage of the genome undergoing each subcompartment transition from proliferating (PRO, columns) to RS (rows); Subcompartments range from A.1.1 (most active) to B.2.2 (most repressed); the lower-right quadrant captures transitions toward more active states. (**D**) Differential expression status (RS vs. PRO) of genes stratified by compartment state in RS: stable B, switching toward B, switching toward A, and stable A. Numbers indicate gene counts in each category. (**E**) Over-representation of curated senescence-associated gene sets among genes switching compartment or subcompartment toward A or toward B in RS. Color reflects log odds ratio; asterisks denote significance (Fisher’s exact test, BH-adjusted; ** FDR < 0.01, *** FDR < 0.001). (**F, G**) Gene ontology (GO) over-representation analysis of genes switching toward the A compartment (F) or toward more active subcompartments (G) in RS. Bars show the top enriched terms ranked by fold enrichment; color indicates –log10 adjusted p-value.

Despite the directional consistency between RS and OIS, the specific genes undergoing compartment switching were largely distinct between the two conditions (Figure 1B). Of the 1,894 genes shifting toward A in RS and the 2,096 in OIS, only 350 were shared; toward-B switching showed even less overlap (34 of 1,079 RS genes and 468 OIS genes). Although these overlaps were statistically greater than expected by chance (Fisher’s exact test, p < 0.01), the predominantly non-overlapping gene sets indicate that, while RS and OIS converge on a global loss-of-heterochromatin trend, the specific genomic regions affected differ substantially between the two senescence programs.

We next integrated our compartment calls with total RNA-seq data, which independently confirmed established senescence-associated transcriptional signatures (Figure S3). Although only a minority of compartment-switching genes were differentially expressed, expression changes were modestly biased in the direction predicted by compartment switching (Figure 1D). Of the 1,894 genes shifting toward A in RS, 437 were significantly upregulated (concordant) versus 112 downregulated (discordant); of the 1,079 genes shifting toward B, 213 were downregulated (concordant) versus 150 upregulated (discordant). Compartment switching is therefore associated with the expected direction of transcriptional change but is not, on its own, a strong predictor of differential expression, consistent with the view that A/B compartmentalization sets a permissive context for transcription but does not by itself determine gene activity.

To resolve compartmentalization at finer scale, we used Calder 2 (Liu et al. 2021) to assign eight subcompartments ranging from A.1.1 (most transcriptionally active) to B.2.2 (most repressed). Subcompartment switching reinforced the loss-of-heterochromatin pattern observed at the compartment level (Figure 1C). The single most frequent transition involved 5.2% of the genome moving from A.1.2 to A.1.1. More broadly, transitions toward more active subcompartments (below the diagonal in the heatmap) substantially outweighed transitions toward more repressive states.

We then asked whether compartment– and subcompartment-level reorganization preferentially affected genes associated with hallmark features of senescence. Among six curated senescence-relevant gene sets tested, only the Reactome Cell Cycle gene set showed significant enrichment (Figure 1E). Cell cycle genes were depleted from regions switching toward A (at both the compartment and subcompartment level) and enriched among regions switching toward B at the subcompartment level (Fisher’s exact test, p ≤ 0.05), consistent with the cell cycle arrest that defines senescence. Notably, canonical inflammatory and SASP-associated programs, including the Hallmark Interferon Alpha Response and the SenMayo gene set, were not significantly enriched among compartment-switching regions, suggesting that activation of these programs in RS is not driven primarily by large-scale compartmentalization changes.

Unbiased gene ontology (GO) analysis of regions shifting toward more active states revealed several enriched processes (Figure 1F, G). Genes moving toward the A compartment were enriched for terms related to complement activation and immune function, including regulation of complement activation (classical pathway), immunoglobulin complex, and adaptive immune response (Figure 1F). At the subcompartment level, the top enriched terms instead reflected cytoskeletal reorganization, intermediate filament organization and intermediate filament-based process, alongside additional immune-related terms (Figure 1G). These enrichments indicate that the loss of heterochromatin in RS exposes a heterogeneous set of loci, including some immune-related genes, but the curated gene-set analysis above argues against coordinated SASP activation through compartment reorganization alone.

### Loop Expansion in RS is Associated with Loss of Methylation and Neuronal Identity Drift

The high resolution of our Hi-C data enabled detailed analysis of chromatin loops in RS, features that require sub-10-kb resolution to call reliably and have therefore been underexplored in prior senescence studies. Loop analysis was not performed on the OIS dataset, as its lower resolution precluded direct comparison with our maps.

To compare loops across conditions, we adopted a loop-matching strategy that classified each loop as condition-specific or shared. We used a window of ±n around each anchor base and called two loops the same when both cross-condition anchor pairs could be found within such a window (Figure S4A). As we swept n from 10 kb to 100 kb on chromosome 21, the counts of unique and shared loops plateaued at ∼30 kb, which we adopted as the matching threshold (Figure S4B); illustrative classifications at 10 kb and 100 kb are shown for reference (Figure S4C). Applying this approach genome-wide, we found 7,623 loops shared between proliferating and RS cells (ps-loops), 1,531 unique to proliferating cells (p-loops), and 8,880 unique to RS cells (s-loops), an approximately sixfold excess of RS-unique over proliferation-unique loops (Figure 2A). RS-unique loops were modestly but significantly longer than both shared loops and proliferation-unique loops (median 300 kb for shared, 340 kb for proliferation-unique, and 370 kb for RS-unique loops; Wilcoxon rank-sum test, BH-adjusted p ≤ 0.001 for both comparisons; Figure 2B), whereas proliferation-unique and shared loops did not differ significantly (p = 0.22).

**Figure 2.**
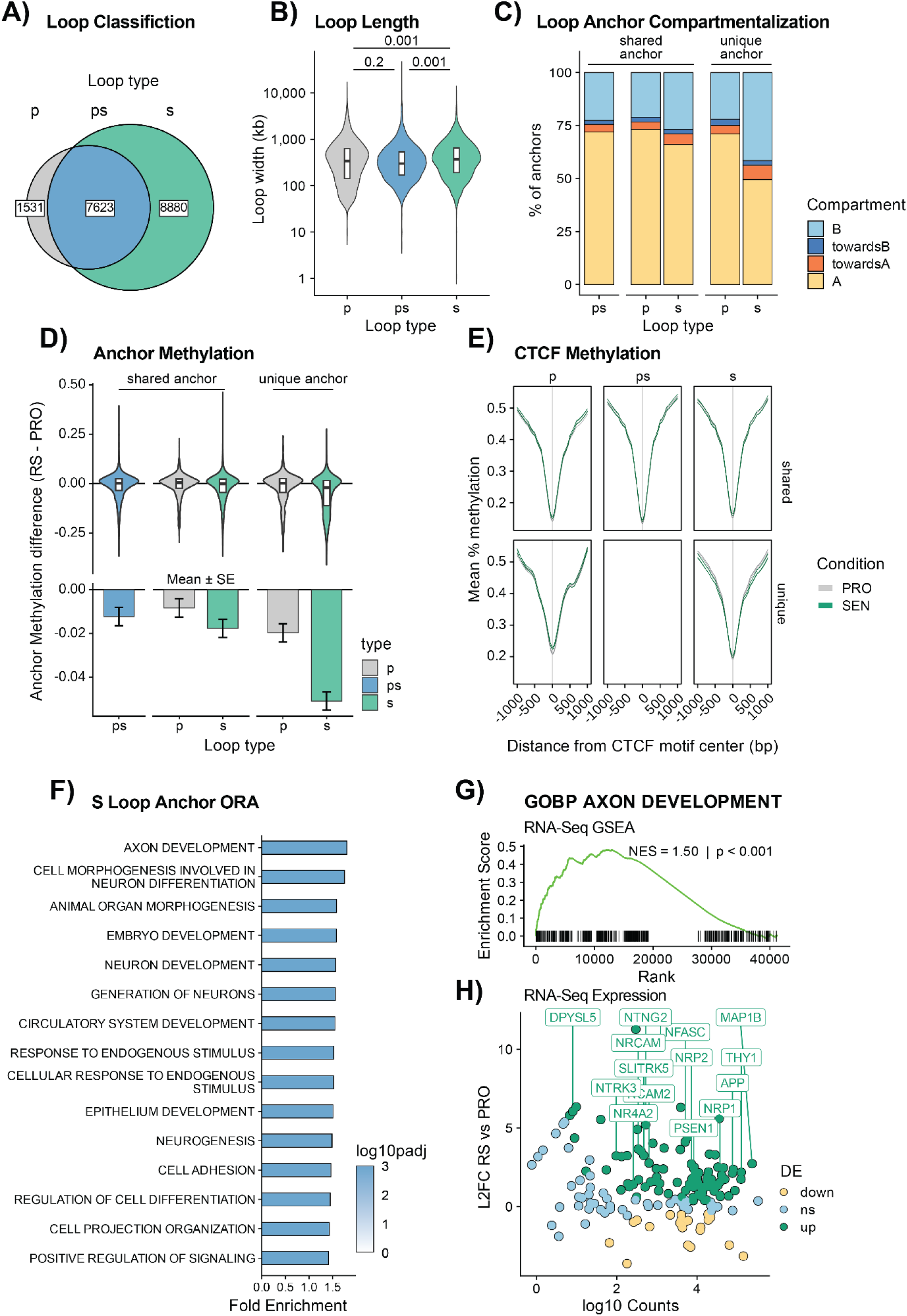
| Loop expansion in replicative senescence is associated with DNA hypomethylation and a neuronal identity-drift gene program. (A) Venn diagram of chromatin loops identified in proliferating (p) and replicatively senescent (s) cells, showing 1,531 loops unique to proliferating cells, 8,880 unique to RS, and 7,623 shared (ps) between conditions. Loops were classified as shared if both anchors fell within ±30 kb of one another (Figure S4). (**B**) Distribution of loop widths (anchor-to-anchor genomic distance) for each loop class. (**C**) Compartment status of loop anchors stratified by whether each anchor was shared or unique to loops of a particular condition. (**D**) CpG methylation difference (RS – PRO) at loop anchors, by anchor class. Top: per-anchor distributions. Bottom: linear model-estimated methylation difference (RS − PRO) ± SE. (**E**) Mean CpG methylation around CTCF site centers (±1 kb) within shared (top) and condition-unique (bottom) loop anchors in proliferating (gray) and RS (green) cells. (**F**) Gene ontology over-representation analysis of genes with promoters (TSS ±5 kb) overlapping s-loop anchors. Color indicates –log10 adjusted p-value. (**G**) RNA-seq gene set enrichment analysis (RS vs. PRO) for the GOBP Axon Development gene set (NES = 1.50, p < 0.001). (H) MA-style plot of Axon Development genes with s-loop anchor promoter overlap, showing log2 fold change (RS vs. PRO) versus log10 mean expression; color indicates differential expression status. Selected upregulated genes are labeled.

We next characterized the genomic context of loop anchors. As with loops, we classified loop anchors as unique or shared. While shared ps-loops by definition have shared anchors, unique loops can be further distinguished by whether both anchors are unique (not present in any loop in the other condition), only one is unique, or both are shared but their pairing is unique. Unique anchors of RS loops (s-anchors) were enriched in the B compartment, and correspondingly depleted from the A compartment, relative to unique p-loop anchors (p-anchors) or shared anchors (Figure 2C; Fisher’s exact test, p ≤ 0.001 for all comparisons). Unique s-anchors were also disproportionately located in regions undergoing compartment switching toward A (Fisher’s exact test, p ≤ 0.001 for all comparisons). In agreement with these findings, the fraction of the B compartment engaged in any loop was substantially higher in RS than in proliferating cells (Figure S5A), indicating that s-loops extend into chromatin territories that are largely loop-poor in proliferating fibroblasts. Most loop anchors overlapped known CTCF sites across all classes, though this proportion was lower for unique anchors than for shared anchors (Figure S5B; Fisher’s exact test, BH-adjusted p ≤ 0.001 for all unique vs. shared comparisons). Consistent with a regulatory role for these architectural changes, genes which gained or lost loop contacts between conditions were more likely to be differentially expressed than genes with stable loop status (Figure S5C; Fisher’s exact test, p ≤ 0.001 for all comparisons).

To investigate why RS cells form so many additional loops, we considered DNA methylation. Genome-wide methylation loss is a hallmark of senescence (Cruickshanks et al. 2013), and DNA hypomethylation has been implicated in chromatin loop formation in other contexts (Zhang et al. 2020). We performed Nanopore long-read sequencing of proliferating and RS LF1 cells in quadruplicate (mean coverage ∼28x per sample, N50 ∼20 kb), enabling accurate genome-wide CpG methylation calling. S-anchors showed substantially greater methylation loss in RS than did p– or shared anchors, with a mean methylation decrease roughly three-fold larger than that of any other anchor class (Figure 2D; linear model with emmeans contrasts, s vs ps difference-of-differences p ≤ 0.001; for all other contrasts, p > 0.05). Unlike 5mC, 5-hydroxymethylcytosine (5hmC), the obligate intermediate of TET-mediated oxidative demethylation, showed no increase at s-anchors relative to other anchor classes in RS (Figure S5E), suggesting that the hypomethylation observed results from either passive, replication-coupled dilution preceding arrest or active demethylation that had already run to completion over the months these cells spent in senescence. Overall, these findings are consistent with a model in which hypomethylation in B-compartment chromatin, together with the broader shift toward A, creates a permissive context for new loop formation.

Because CTCF binding is sensitive to CpG methylation within its conserved motif (H. Wang et al. 2012; Hashimoto et al. 2017), we asked if RS-unique loop anchor CTCF sites exhibited changes in methylation. Methylation profiles around CTCF sites within loop anchors showed the canonical “valley” centered on the motif but were essentially superimposable between proliferating and RS cells across all anchor classes, including s-anchors (Figure 2E). Thus, although the broader s-anchor regions lose methylation in senescence, the CTCF sites themselves do not, arguing that altered CTCF binding through direct methyl-CpG remodeling at the CTCF site is unlikely to be the proximate driver of the senescence-associated loop expansion.

We next asked whether gene programs were physically associated with s-loops. Because s-loops cover >80% of the genome (Figure S5A) these loops cover the majority of genes in the majority of gene sets. We therefore focused on the more limited set of genes whose promoters (TSS ±5 kb) overlap s-loop anchors, rather than those within the loop interior. This choice is supported by prior work showing that gained loops drive concordant changes in gene expression at genes whose promoters overlap loop anchors, but not at genes between anchors (Bond et al. 2023; Reed et al. 2022). Gene ontology analysis revealed enrichment for neuronal differentiation and morphogenesis terms: the top hit was Axon Development, and other enriched terms included Cell Morphogenesis Involved in Neuron Differentiation, Neuron Development, Generation of Neurons, and Neurogenesis (Figure 2F). Although fibroblasts and neurons arise from distinct germ layers, the two cell types lie close in the epigenetic landscape: forced expression of just three transcription factors (Ascl1, Brn2, and Myt1l) is sufficient to directly convert fibroblasts into functional neurons without passage through a pluripotent intermediate (Wapinski et al. 2013; Vierbuchen et al. 2010). Our finding that s-loops mark neuronal-lineage loci therefore identifies senescence as a state in which this latent epigenetic proximity may be partially realized at the level of 3D contacts. Consistent with this interpretation, Axon Development gene set was significantly enriched among genes upregulated in RS by RNA-seq (NES = 1.50, p < 0.001; Figure 2G), and many Axon Development genes with promoters overlapping s-loop anchors were significantly upregulated (Figure 2H).

Together, these results show that senescence-associated loop expansion is enriched in the B compartment, is accompanied by anchor-region DNA hypomethylation that does not extend to CTCF sites themselves, and is associated with an upregulation of neuronal-lineage genes. We interpret this as a structural correlate of the loss of cell-type identity that has recently been described as a feature of aging chromatin (Yang et al. 2023), with senescent fibroblasts ectopically activating lineage-inappropriate gene programs in regions of newly formed 3D contacts.

### Retrotransposon Expression is Associated with Compartment and Subcompartment Status

Senescence is accompanied by widespread upregulation of retrotransposable elements, which can drive sterile inflammation in aging tissues (De Cecco, Criscione, Peckham, et al. 2013; De Cecco, Criscione, Peterson, et al. 2013). In particular, derepression of L1 elements activates the cytosolic DNA–sensing pathway and contributes to interferon-driven SASP induction (De Cecco et al. 2019). Whether the 3D genome reorganization we observe in RS contributes mechanistically to RTE derepression has not previously been examined. To address this, we integrated our compartment, subcompartment, and loop calls with locus-resolved RTE expression estimates. Locus-level quantification of RTE expression is challenging because of high inter-element sequence similarity and the presence of polymorphic, non-reference insertions absent from standard reference assemblies. To improve short-read mapping accuracy, we generated a custom LF1 reference genome that augments the T2T-CHM13 reference genome with non-reference L1 insertions identified from deep Nanopore long-read sequencing and masks absent reference elements (see Methods). We then used Telescope, which applies an expectation-maximization algorithm to assign multi-mapping reads to their most probable locus. To prevent overlapping gene transcription from confounding our RTE expression estimates, we restricted our analysis to intergenic and genic antisense RTEs.

Consistent with prior reports, we observed broad family-level upregulation of full-length RTEs in RS (Figure 3A, top; Figure S3E). To assess the contribution of chromatin context to these transcriptional changes, we compared the compartment distribution of each family’s genomic copies to the compartment distribution of its expressed RNA (Figure 3A, bottom). Across nearly all families, A and toward-A compartments contributed disproportionately to expression relative to their genomic occupancy, while B-compartment copies contributed proportionally less. This pattern indicates that, at the family level, RTE expression in RS is biased toward copies residing in or moving into active compartments, rather than being uniformly distributed across the genome.

**Figure 3.**
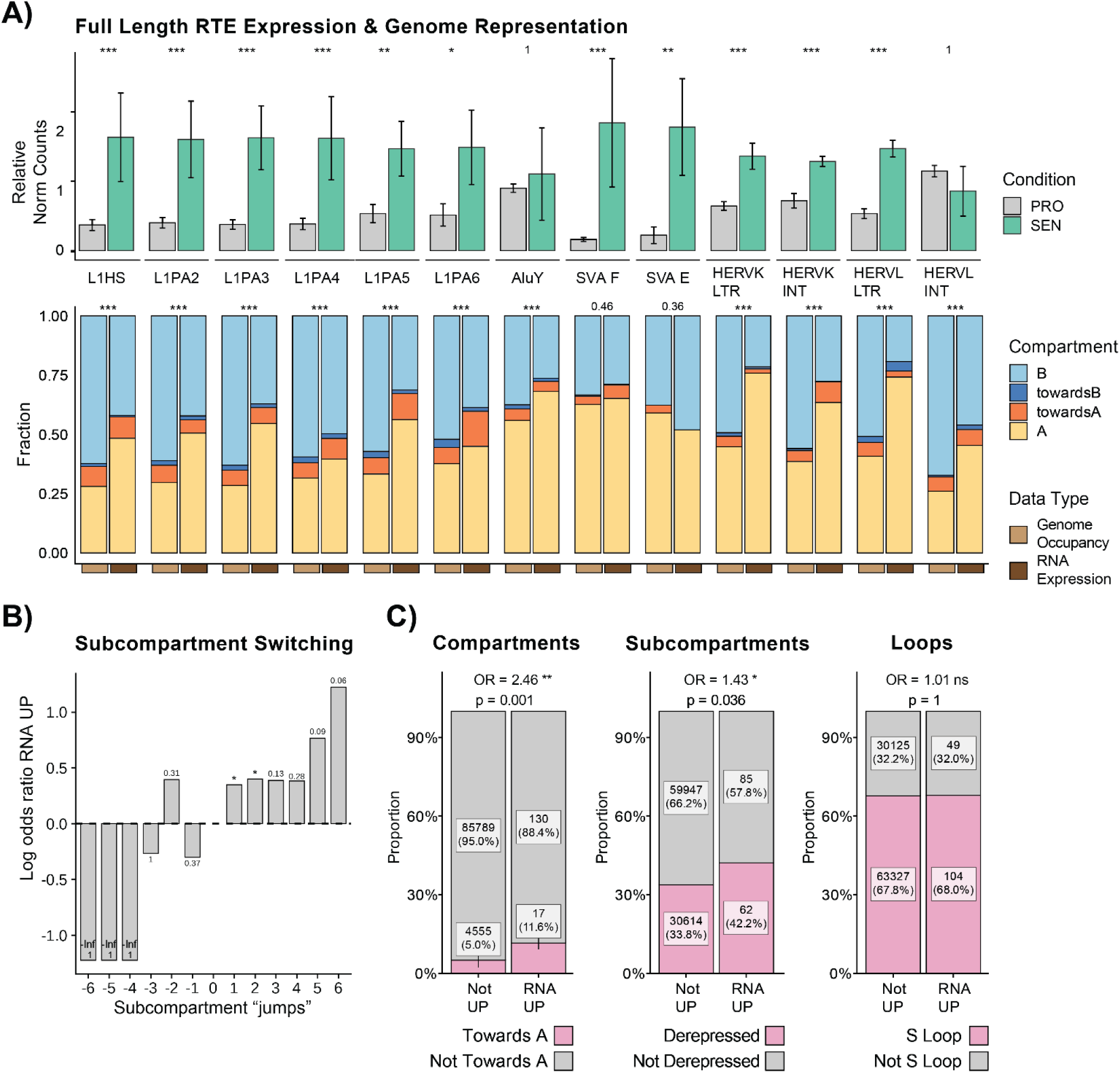
| Senescence-associated compartment and subcompartment reorganization, but not loop status, is associated with RTE derepression. To avoid bias from overlapping gene transcription, RTE expression analyses were restricted to intergenic and genic antisense elements. (A) Top: Family-level normalized expression of full-length retrotransposable elements (RTEs) in proliferating (PRO) and replicatively senescent (RS) cells. Counts were normalized using DESeq2 size-factors and scaled relative to each subfamily’s mean expression across all samples. Bars show mean ± SE across replicates. Statistical significance was assessed using a negative binomial GLM incorporating DESeq2-derived size factors as a model offset; asterisks denote significance (* FDR ≤ 0.05, ** FDR ≤ 0.01, *** FDR ≤ 0.001). Bottom: Compartment distribution of full-length members of selected RTE subfamilies in RS, comparing the genomic occupancy of each family (left bar) to the share of family-level RNA expression originating from each compartment (right bar). For most families, A and toward-A compartments contribute more to expression than their genome occupancy would predict. Statistical significance was assessed using a chi-squared goodness-of-fit test comparing each replicate’s expression compartment distribution to the genomic element proportions; the maximum BH-adjusted p-value across replicates is shown for each subfamily; asterisks denote significance (*** FDR ≤ 0.001). (B) Log odds ratio of being upregulated in RS for individual RTE loci as a function of subcompartment “jumps” between proliferating and RS cells. Each bar represents a cumulative threshold: at jump = k, the odds ratio compares loci that jumped at least k subcompartments toward an active (positive k) or repressed (negative k) state against all other loci. Numbers next to each bar are unadjusted two-sided Fisher’s exact test p-values; because thresholds are cumulative, the tests are nested and not independent; asterisks denote significance (* p ≤ 0.05). (C) Proportion of individual RTE loci classified as significantly upregulated in RS (RNA UP) versus not, stratified by (left) compartment switching toward A, (middle) subcompartment switching toward a more active state (“derepressed”), and (right) presence within an RS-specific loop. Odds ratios and Fisher’s exact test p-values are shown above each panel.

We next asked whether subcompartment switching predicted locus-specific upregulation. We stratified full-length RTEs by the number of subcompartment “jumps” between proliferating and RS cells, from –6 (most repressed) to +6 (most activated). Loci that shifted toward more active subcompartments were modestly more likely to be upregulated (Figure 3B): the odds of upregulation were elevated among loci with at least one activating jump (OR = 1.42, Fisher’s exact test p = 0.045) and at least two activating jumps (OR = 1.49, p = 0.046). Odds ratios remained above 1 for larger activating shifts (OR = 1.5–3.4 for ≥ +3 to ≥ +6 jumps) but did not reach significance, as these categories contained few upregulated loci (3–17). Loci shifting toward more repressed subcompartments showed no enrichment for upregulation.

We then asked whether each architectural feature, compartment switching, subcompartment switching, or loop membership, was independently associated with RTE derepression. Compartment switching toward-A was the strongest predictor: 11.6% of upregulated RTE loci resided in toward-A regions, compared to 5.0% of non-upregulated loci (OR = 2.46, Fisher’s exact test p = 0.002; Figure 3C, left). Subcompartment switching toward more active states was also significantly associated with upregulation, though more modestly: 42.2% of upregulated loci were in derepressed regions versus 33.8% of non-upregulated loci (OR = 1.43, Fisher’s exact test p = 0.036; Figure 3C, middle). In contrast, residence within an s-loop showed no association with RTE derepression: 68.0% of upregulated loci and 67.8% of non-upregulated loci were within s-loops (OR = 1.01, Fisher’s exact test p = 1; Figure 3C, right).

Together, these results indicate that compartment– and subcompartment-level reorganization, but not the senescence-associated loop expansion described above, is associated with RTE derepression in RS. This is consistent with a model in which the loss of heterochromatin documented in Figure 1 creates a more permissive environment for RTE transcription.

### Chromatin Remodeling Across Multiple Architectural Scales Converges on L1HS_14q23.2_3, a Derepressed Intact L1HS Element Which Loses Promoter Methylation in RS

We focused subsequent analysis on intact L1HS elements, which are the only autonomously retrotransposition-competent elements in the human genome and the dominant endogenous source of active reverse transcriptase (RT) activity (Beck et al. 2010; Brouha et al. 2003). Aggregated across all intact, full-length L1HS elements (those retaining open reading frames for both ORF1p and ORF2p), expression was significantly elevated in RS relative to proliferating cells (Figure 4A; negative binomial modeling of counts, incorporating DESeq2-derived size factors as a model offset, FDR = 0.02). At the level of individual loci, however, the picture was sparser: of the intact L1HS elements detectable by RNA-seq, only two were significantly upregulated in RS (FDR ≤ 0.05): L1HS_14q23.2_3, an intergenic insertion, and L1HS_7p14.3_1, a genic antisense insertion (Figure 4B). The small number of individually significant loci likely stems from (i) the inherent statistical limitations of assigning multi-mapping, short-read signal to highly similar elements at low per-locus coverage (which can lead to false positive and negative expression calls), and/or (ii) genuine locus-specific derepression.

**Figure 4.**
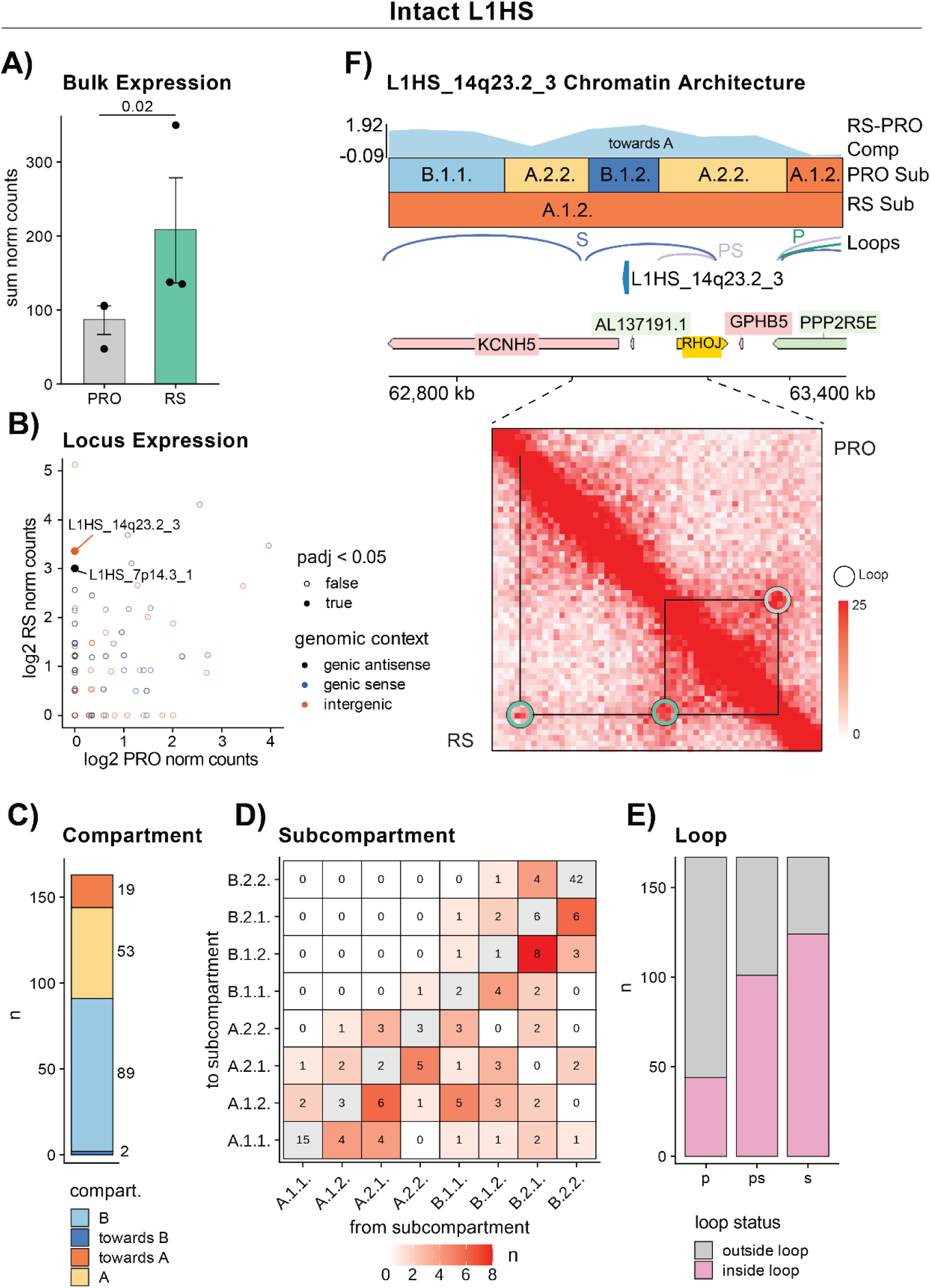
| Chromatin remodeling at multiple architectural scales is associated with the derepression of an intact L1HS element. (**A**) Sum of normalized expression counts across intact L1HS elements in proliferating (PRO) and replicatively senescent (RS) cells. Bars show mean ± SE across replicates. Statistical significance was assessed using a negative binomial GLM incorporating DESeq2-derived size factors as a model offset; FDR = 0.02. (**B**) Per-locus expression of intact L1HS elements in PRO versus RS (log2 normalized counts), colored by genomic context (intergenic, genic sense, genic antisense). Filled points indicate loci with significant differential expression (FDR ≤ 0.05). The two significantly upregulated loci, intergenic L1HS_14q23.2_3 and genic antisense L1HS_7p14.3_1, are labeled. (**C**) Compartment distribution of intact L1HS elements in RS cells. (**D**) Subcompartment distribution of intact L1HS elements, plotted as a heatmap of transitions from PRO (columns) to RS (rows). (**E**) Number of intact L1HS elements located inside (purple) or outside (gray) each loop class. (**F**) Local chromatin architecture surrounding L1HS_14q23.2_3 on chromosome 14. Top: dcHi-C track (RS – PRO compartment score), subcompartment calls in PRO and RS, loops, L1HS_14q23.2_3 position, and gene annotations (GTF). Bottom: Hi-C contact map of the region, with PRO above the diagonal and RS below; circles mark chromatin loops. At 5-kb resolution, this loop is called by HiCCUPS in RS (observed = 26; 4.5-fold enrichment over the local donut background; significant against all four HiCCUPS background models, q_donut < 0.0001) but is not called in PRO, where both anchors nonetheless anchor other significant loops (q ≤ 0.02), indicating an RS-specific loop gain rather than a difference in mappability or coverage.

To help disambiguate the contributions of these two scenarios, we examined chromatin architectural features of intact L1HS elements. At the compartment level, most intact L1HS loci resided in stable B or A regions, with smaller fractions undergoing compartment switching (Figure 4C). At the subcompartment level, while transitions toward more active states predominated, the largest single group of elements remained in B.2.2 → B.2.2 (no change but heavily repressed; n = 42), with a smaller subset shifting toward more permissive states (Figure 4D). With respect to looping, intact L1HS elements were broadly distributed inside and outside each loop class, with the largest absolute number falling within s-loops (Figure 4E). The architectural context of intact L1HS elements is therefore highly heterogeneous, providing a basis on which 3D-genome structure can confer locus-specific transcriptional advantages/disadvantages to elements which carry highly similar 5′UTR promoter sequences.

We next focused on the chromatin architectural and epigenetic features of the two significantly upregulated intact L1HS loci. L1HS_7p14.3_1 was located in stable A-compartment chromatin and shifted only modestly within A (A.2.1 → A.1.2), and hence exhibited no marked compartment– or subcompartment-level reorganization to distinguish it from the bulk of intact L1HS elements. Although it fell within an s-loop, its host gene, NOD1, was downregulated in senescence. The L1HS_7p14.3_1 5′UTR (the L1 internal promoter region) was highly methylated in both proliferating and senescent cells. These data suggest that L1HS_7p14.3_1 upregulation may be spurious, an artifact of multi-mapping read reassignment, a known limitation of this method. By contrast, at L1HS_14q23.2_3, derepressive architectural changes converged across all scales (Figure 4F): a marked toward-A compartment shift, a B.1.2 → A.1.2 subcompartment transition (one of the most pronounced activating jumps in the heatmap), and a new s-loop bringing it into contact with two neighboring genes, KCNH5 and RHOJ, both upregulated in RS and the latter particularly highly expressed (Figure S5D). The Hi-C contact map showed this loop present in the RS map (below diagonal) but absent from proliferating cells (above diagonal). Methylation across the element’s body and surrounding ±10 kb was reduced in RS (Figure 5A), but the most important change occurred over the first ∼8 CpGs of its 5′UTR, where roughly half of the sequencing reads in RS showed near-complete loss of methylation while nearly all reads in proliferating cells remained fully methylated (Figure 5B). Quantitatively, L1HS_14q23.2_3 ranked at the >95th percentile among full-length L1HS elements for promoter methylation loss across these eight CpGs (in contrast, L1HS_7p14.3_1 was at the 22nd percentile).

**Figure 5.**
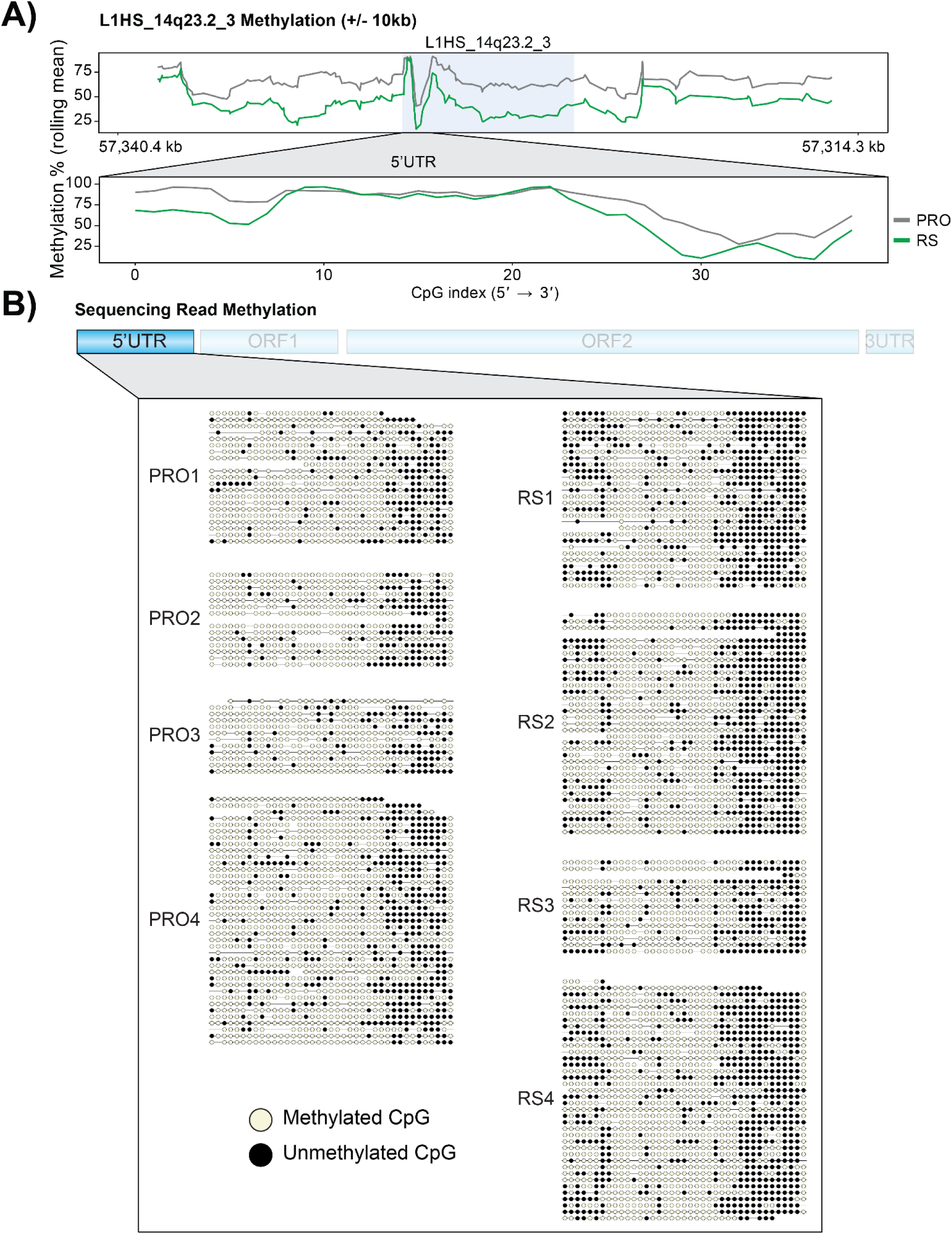
| L1HS_14q23.2_3 undergoes pronounced promoter hypomethylation in replicative senescence. (**A**) Average CpG methylation (rolling mean over Nanopore reads, k = 3) across the L1HS_14q23.2_3 locus ±10 kb in proliferating (PRO, gray) and RS (green) cells. L1HS_14q23.2_3 is highlighted in gray. Below: zoomed view of the 5′UTR, showing per-CpG methylation indexed 5′→3′ across the L1 promoter. (**B**) Single-read methylation maps spanning the L1HS_14q23.2_3 5′UTR for each Nanopore replicate (PRO1–4, left; RS1–4, right). Each row is one read; each circle is one CpG. Filled circles, unmethylated; open circles, methylated.

Taken together, several architectural and epigenetic changes converge at L1HS_14q23.2_3: a compartment-level shift toward A, a strong activating subcompartment transition, the formation of a new senescence-specific loop topologically linking it to two upregulated genes, and pronounced promoter demethylation. This same locus emerges as one of a small handful of significantly upregulated intact L1HS elements across two additional senescence datasets from our group: DNA-damage-induced senescence in human astrocytes (Woodham et al. 2026) and a recently published LF1 replicative senescence dataset in which L1HS_14q23.2_3 was upregulated in the transcriptome and translatome (the translationally engaged pool of mRNA) (Kelsey et al. 2026). The recurrence of L1HS_14q23.2_3 derepression across distinct senescence-inducing stimuli and cell types favors a model in which the population of RT-encoding L1HS elements actively transcribed in senescent cells may be small and architecturally selected, with L1HS_14q23.2_3 as a leading candidate.

### Quiescent Cells Display Senescence-Associated Inflammatory Features and are not a Suitable Senescence Control

Beyond proliferating and RS cells, we generated RNA-seq data and a 2.5 kb-resolution Hi-C map for quiescent (QUI) LF1 fibroblasts, induced by contact inhibition followed by serum withdrawal. Quiescence was originally selected as a control for senescence because it represents a more physiological, reversible cell cycle arrest commonly observed in fibroblasts within tissues. Gene expression principal component analysis (PCA) showed QUI separating cleanly from both PRO and RS along orthogonal axes: PC1 (66% of variance) distinguished arrested from proliferating cells, while PC2 (27%) separated QUI from RS (Figure 6A). As expected, QUI showed pronounced downregulation of Hallmark E2F Targets (Figure 6B) and uniquely upregulated CDKN1C (Figure 6E), confirming that the cells had exited the cell cycle. However, a more comprehensive Hallmark gene set enrichment analysis (GSEA) of QUI versus PRO revealed an unexpectedly senescence-like transcriptional profile (Figure 6C). Alongside the expected downregulation of proliferation-associated programs (E2F Targets, MYC Targets V1/V2, G2M Checkpoint), QUI cells showed strong upregulation of programs typically associated with senescence and the SASP, including Interferon Alpha Response, TNFA Signaling via NFκB, and Inflammatory Response.

**Figure 6.**
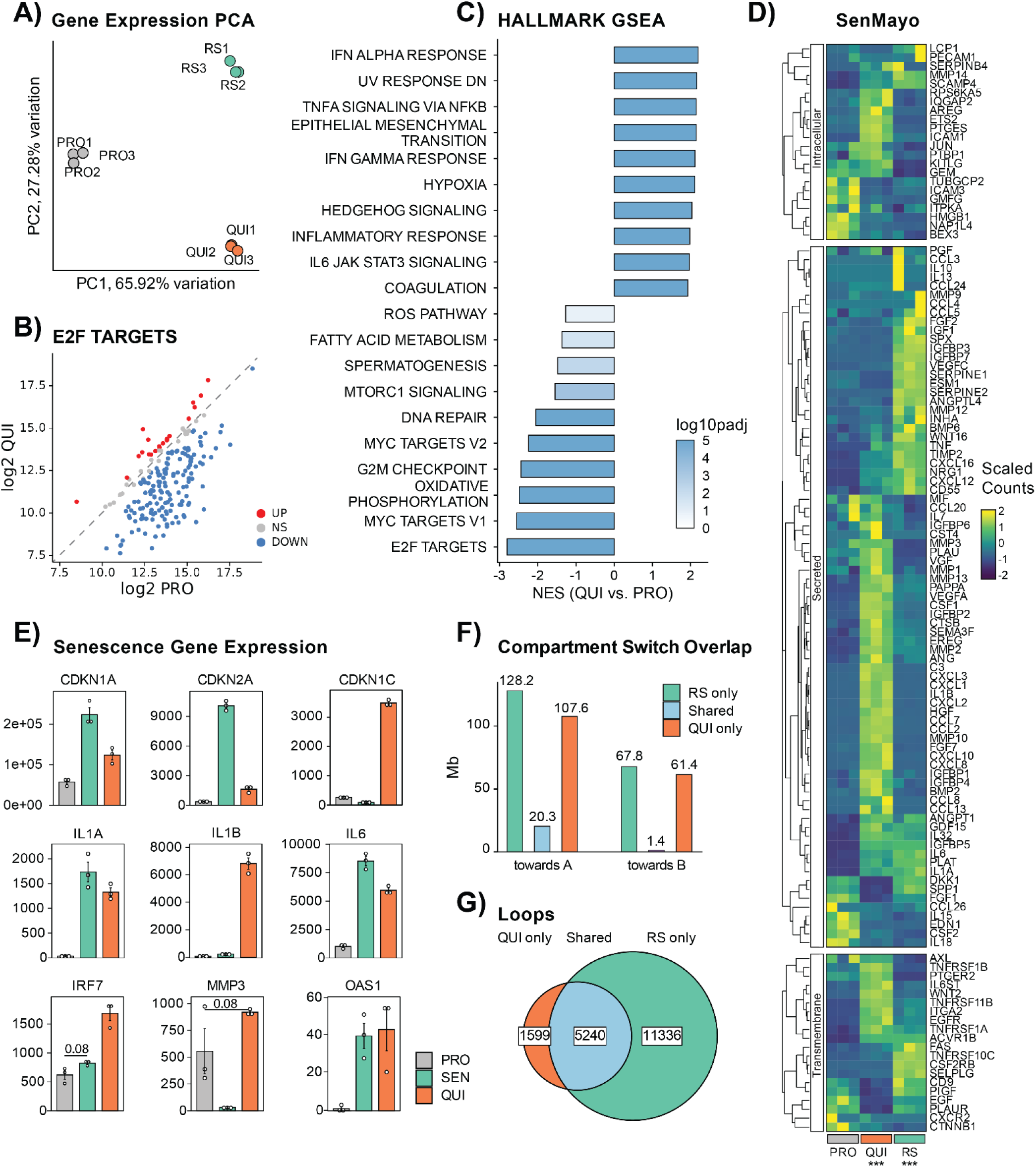
| Quiescent fibroblasts share inflammatory and senescence-associated transcriptional features but show distinct chromatin architectural changes. (A) Principal component analysis of total RNA-seq data from proliferating (PRO), quiescent (QUI), and replicatively senescent (RS) LF1 fibroblasts. PC1 (65.92% variation) separates arrested from proliferating cells; PC2 (27.28% variation) separates QUI from RS. (B) Hallmark E2F Targets gene expression in QUI versus PRO. Genes are colored by differential expression status (DOWN, NS, UP). (C) Hallmark GSEA of QUI versus PRO. Bars show normalized enrichment scores (NES); color reflects –log10 adjusted p-value. (D) Heatmap of SenMayo gene expression across PRO, QUI, and RS replicates, separated into intracellular, secreted, and transmembrane subsets. Counts are scaled per gene. Asterisks below QUI and RS indicate significant enrichment of the SenMayo signature relative to PRO by GSEA (*** FDR < 0.001). (E) Expression of representative senescence and SASP genes across PRO, RS, and QUI from total RNA-seq. Bars show mean ± SE. SEN and QUI were compared to PRO and for all genes, these comparisons were significant (FDR ≤ 0.05) except for those with p-values shown. (F) Megabases of the genome undergoing compartment switching toward A or toward B (vs. PRO) in RS-only, QUI-only, or both. (G) Venn diagram of chromatin loops identified in QUI and RS, showing 1,599 QUI-only, 11,336 RS-only, and 5,240 shared loops.

The SenMayo gene set, a curated senescence signature largely composed of SASP factors, was significantly enriched in both QUI and RS relative to PRO (Figure 6D). The two arrested states activated shared and distinct subsets of SenMayo (Figure 6D). Direct inspection of canonical senescence and SASP transcripts confirmed this pattern (Figure 6E). The cell cycle inhibitors CDKN1A (p21) and CDKN2A (p16) were both substantially upregulated in QUI relative to PRO, although less so than in RS. The cytokine IL1B and the interferon regulator IRF7 were strongly upregulated in QUI, above RS levels. We did not observe increased expression of FL L1HS elements in QUI relative to PRO (Figure S6A) which suggests that these RTEs are not the cause of the elevated inflammatory signalling observed in QUI.

To test whether this inflammatory phenotype was an artifact of our particular induction protocol, we performed RT-qPCR on a panel of senescence and SASP genes in IMR-90 fibroblasts, a fetal lung fibroblast strain closely comparable to LF1, subjected to contact inhibition (CI) or serum withdrawal (SW) (Figure S6B). Both methods led to upregulation of SASP genes and classical senescence markers such as CDKN1A (p21) and CDKN2A (p16), further supporting the inflammatory nature of these commonly used in vitro quiescence models.

Together, these results indicate that quiescent fibroblasts induced by standard in vitro protocols share a substantial fraction of the senescence-associated inflammatory transcriptional program, including upregulation of SenMayo genes, classical senescence markers, and SASP cytokines. Using such cells as a senescence control would therefore mask the inflammatory features of senescence relevant to aging biology. We accordingly chose to analyze QUI as a separate condition rather than as a senescence reference.

We compared chromatin architecture between QUI and RS at the same scales examined in earlier sections. At the compartment level, QUI and RS showed similar overall fractions of the genome shifting toward A and toward B (vs. PRO), but the specific regions involved were largely distinct (Figure 6F). Of the megabases switching toward A, 128.2 Mb were RS-only, 107.6 Mb were QUI-only, and only 20.3 Mb were shared. The asymmetry was even more pronounced for switching toward B, where 67.8 Mb were RS-only and 61.4 Mb were QUI-only, but only 1.4 Mb were shared. ChromHMM analysis of these regions (Figure S6D) showed broadly similar regulatory state composition across RS-only and QUI-only regions. At the subcompartment level, QUI exhibited a different switching pattern from RS (Figure S6C). Whereas RS was dominated by activating transitions (A.1.2 → A.1.1 at 5.2%, Figure 1C), the most frequent transition in QUI was the inverse (A.1.1 → A.1.2 at 4.7%), and QUI showed no overall preference for activating versus repressing shifts. At the loop level, we identified 6,839 loops in QUI cells. Comparing QUI and RS, we found 1,599 QUI-only loops, 11,336 RS-only, and 5,240 shared (Figure 6G). The 7-fold excess of s-loops over q-loops parallels the loop expansion observed in the PRO-versus-RS comparison (Figure 2A) and indicates that the increase in chromatin loops is specific to senescence rather than a general feature of cell cycle arrest.

In summary, although quiescence shares features of senescence at the level of compartment shift magnitude and SASP gene activation, the specific chromatin regions undergoing reorganization differ markedly between the two states, the subcompartment switching patterns are not shared, and the loop expansion seen in RS is not recapitulated. These results constitute the first multi-scale architectural comparison between the two arrested states.

## Discussion

The interplay between three-dimensional genome organization and gene regulation has emerged as a central theme in development, disease, and aging (Krumm and Duan 2019). In parallel, cellular senescence has gained recognition as a driver of age-associated dysfunction, primarily through its inflammatory secretory phenotype, fueled in part by the derepression of retrotransposable elements that activate cytosolic nucleic acid sensors (De Cecco et al. 2019; Suryadevara et al. 2024; B. Wang et al. 2024). Here, we integrated the highest-resolution Hi-C maps of replicatively senescent cells generated to date with total RNA-seq and long-read DNA-seq-derived whole-genome methylation profiles. Three principal observations emerge: a global loss of heterochromatin evident at both the compartment and subcompartment levels; an expansion of chromatin loops concentrated in the B compartment, accompanied by anchor-region DNA hypomethylation and associated with activation of lineage-inappropriate gene programs; and a structural basis for retrotransposon derepression, exemplified by an intact L1HS element that we identify as a recurrent locus of derepression across multiple senescence contexts.

Our compartment-level findings reinforce the longstanding loss-of-heterochromatin model of senescence (Rocha et al. 2022; Criscione, Teo, et al. 2016). Both RS and OIS showed asymmetric switching toward the active A compartment, and even regions already classified as A underwent further activation at the subcompartment level. Although the directionality of these changes was shared between RS and OIS, the specific genomic regions involved were largely distinct, consistent with the broader view that senescence is a family of related arrest programs with stimulus-specific footprints. Compartment switching, however, was only modestly predictive of differential gene expression. The clearest signal among curated senescence-associated gene sets was the depletion of cell-cycle genes from regions switching toward A and their enrichment in regions switching toward repressive subcompartments, consistent with cell-cycle arrest being the feature of the senescent program most directly encoded in genome architecture. Notably, SASP and interferon gene sets were not significantly enriched among compartment-switching regions, arguing that the inflammatory secretome is not directly templated by large-scale heterochromatin loss and is more likely controlled by transcription factor–level regulation. Where compartment-level reorganization does show a clear association, as we develop below, is with retrotransposon derepression.

Senescence-specific chromatin loops were in approximately sixfold excess relative to proliferation-specific loops, a feature resolvable only at the sub-10-kb resolution achieved here. S-loops were concentrated in the B compartment and accompanied by DNA hypomethylation across their anchor regions. Two lines of prior work connect DNA methylation and cohesin dynamics to three-dimensional genome folding, and our findings are distinct from both. Zhang and colleagues showed that large hypomethylated nadirs anchor very long-range, often inter-chromosomal contacts spanning many megabases, but these formed independently of CTCF– and cohesin-mediated extrusion and their anchors lacked CTCF (Zhang et al. 2020). The senescence-specific loops we describe differ in scale (∼370 kb versus many megabases), compartment context, and occur at anchors that predominantly overlap known CTCF sites. Separately, Olan et al. found that oncogene-induced senescence rewires loops through cohesin redistribution rather than altered CTCF binding, with de novo cohesin recruited to the 3′ ends of active genes in a transcription-dependent, CTCF-independent manner (Olan et al. 2020). Our data share the broad principle that senescence can reorganize looping without remodeling CTCF motifs – indeed, we find that anchor regions lose methylation while the CTCF motifs within them do not – but point to a different proximate route. Because s-loops are concentrated in the B compartment, we favor a model in which posits that the genome-wide heterochromatin loss and decondensation characteristic of replicative senescence render formerly compact B-compartment territory newly accessible to cohesin-mediated extrusion, permitting loop formation at pre-existing CTCF sites without local CpG remodeling. The B-compartment enrichment of s-loops, the anchor-broad but motif-sparing pattern of methylation loss, and the accompanying shift toward the A compartment are together consistent with this picture.

The gene programs encompassed by these new loops are unexpected. Genes with promoters overlapping s-loop anchors were enriched for neuronal differentiation and morphogenesis terms, with Axon Development as the top hit, and many were concordantly upregulated in RS. Senescent fibroblasts thus appear to ectopically activate a transcriptional program characteristic of an entirely different lineage. This finding aligns with recent observations from the ICE model of aging, in which loss of epigenetic information drives gain of lineage-inappropriate enhancer–promoter contacts and ectopic gene expression (Yang et al. 2023). Our data suggest that senescence, like normal aging, is accompanied by a structural blurring of cell-type identity, with 3D contacts that would normally promote lineage fidelity reorganized to expose alternative gene programs. Whether this neuronal-lineage drift has functional consequences for the senescent cell, contributes to age-related tissue dysfunction in vivo, or simply reflects stochastic loss of fibroblast identity remains an open question.

Our analysis of chromatin architecture at RTE loci provides a structural complement to prior reports of RTE derepression in senescence (De Cecco et al. 2019; Simon et al. 2019). At the family level, full-length L1, Alu, SVA, and HERV elements were upregulated in RS, with elements residing in A and toward-A compartments contributing disproportionately to family-level expression. At the locus level, compartment switching toward A (OR = 2.46) and subcompartment switching toward more active states (OR = 1.43) were both significantly associated with RTE upregulation. In contrast, residence within an s-loop showed no association with RTE upregulation. Thus, the global loss of heterochromatin, not the formation of new chromatin loops, provides an architectural basis for transposon derepression in senescence.

A central challenge in the field has been to identify which specific L1 elements contribute to the inflammatory phenotypes of senescent cells. We identified two intact L1HS elements as significantly upregulated in RS. Of these, L1HS_14q23.2_3 emerged as a structural and epigenetic hotspot at which all four features examined in this study converged: a compartment-level shift toward A; a strong activating B.1.2-to-A.1.2 subcompartment transition; the formation of a senescence-specific loop bringing it into proximity with two upregulated genes (KCNH5 and RHOJ); and demethylation of its 5′UTR promoter. As described in the Results, this same locus also emerged as significantly upregulated in two independent senescence datasets analyzed by our group, spanning a different RS protocol, a distinct senescence-inducing stimulus (DNA-damage-induced), and a different cell type (astrocytes). Together, the architectural and epigenetic convergence at L1HS_14q23.2_3 and its recurrence across senescence contexts identify it as a strong candidate for a structurally selected, retrotransposition-competent L1 putatively contributing cytosolic reverse transcriptase (RT) activity and hence fueling interferon-driven inflammation in senescent cells. Functional dissection of this locus, including direct tests of whether L1HS_14q23.2_3 generates RT-activity capable of triggering cGAS–STING signaling and whether targeted silencing of this locus attenuates the inflammatory secretome, is an important next step.

Our quiescence comparison yields an unexpectedly cautionary result. Quiescent LF1 fibroblasts showed a transcriptional profile that overlapped substantially with senescence: SenMayo enrichment was significant, classical senescence markers including CDKN1A and CDKN2A were upregulated, and many SASP cytokines were elevated. This phenotype was recapitulated by contact inhibition and serum withdrawal experiments in IMR-90, suggesting that it is intrinsic to in vitro quiescence induction rather than to a particular cell strain or protocol. These results suggest that quiescent fibroblasts as currently induced in vitro are therefore not a clean control for senescence and could obscure inflammatory features relevant to aging biology.

A key limitation of this study is that our architectural conclusions rest on deep Hi-C maps from a single cell strain, LF1 fibroblasts. Establishing how broadly the compartment– and loop-level reorganization we describe generalizes will require comparable high resolution maps of replicative senescence in other cell types.

Together, these results provide the highest-resolution view of the senescent 3D genome to date and connect architectural reorganization at multiple scales to two transcriptional features of senescence with direct relevance to aging: the structural exposure of lineage-inappropriate gene programs and the derepression of retrotransposable elements. By identifying L1HS_14q23.2_3 as a recurrently derepressed, structurally and epigenetically remodeled intact L1HS locus, our work provides a concrete, locus-level entry point for testing whether specific endogenous L1 elements drive the chronic inflammation of aged tissues.

## Materials and Methods

### Cell Culture

Female human diploid lung fibroblast cells (LF1) were used for all Hi-C, RNA-seq, and Nanopore long-read DNA sequencing experiments. Cells were maintained in growth media consisting of HAMS F10, 15% FBS, and 1× Penicillin/Streptomycin/Glutamine, at 37°C, 5% CO₂, and 2.5% O₂. Proliferating cells were passaged in regular growth media and harvested at 60–80% confluency. Quiescent cells were obtained by growing them to 100% confluency, then switching to serum-free media for 1–2 days before harvest. Replicatively senescent (RS) cells were generated by serial passaging in regular growth media until replicative exhaustion, then maintained in the senescent state for 3–4 months before harvest to ensure late-stage senescence. The independent IMR-90 RT-qPCR validation experiments used a different culture protocol and quiescence was induced by contact inhibition (CI) or serum withdrawal (SW) instead of a combination of both; see “RT-qPCR Validation in IMR-90 Cells” below.

### Hi-C Library Preparation and Sequencing

For each condition (proliferating, quiescent, RS), three biological replicates were prepared. Cells were crosslinked with 1% paraformaldehyde at room temperature for 10 minutes while attached to cell culture plates. Crosslinking was quenched with glycine. Cells were washed three times with PBS and scraped into microcentrifuge tubes. Hi-C libraries were prepared using the Arima Genomics Hi-C kit according to the manufacturer’s instructions. For each condition, the three libraries were pooled and sequenced across three lanes of an Illumina HiSeq, yielding approximately one billion paired-end reads (Supplementary Table 1).

### Hi-C Read Processing and Map Generation

Raw Hi-C reads from this study, along with the publicly available oncogene-induced senescence (OIS) Hi-C dataset (GEO: GSE135093), were aligned to the hg38 reference genome and assembled into Hi-C contact maps using the Juicer pipeline version 1.6 (Durand et al. 2016). Replicate reproducibility was assessed using HiCRep (Lin et al. 2021) at 100 kb resolution. After confirming reproducibility, replicates were merged within each condition to produce deeply sequenced contact maps; map resolutions were determined using Juicer’s standard MAPQ ≥ 30 read coverage criterion, yielding 3.35 kb for proliferating cells, 2.75 kb for RS cells, and 2.5 kb for quiescent cells. Hi-C maps were visualized using CoolBox (Xu et al. 2021) and Juicebox (Durand et al. 2016). Default Juicer cutoffs (20 kb) were used to define long-range and short-range contacts. t-tests were used to compare the percentage of intra-chromosomal contacts across replicates.

### Compartment (A/B) Analysis

A/B compartments and inter-condition compartment switching were called using dcHi-C (Chakraborty et al. 2022) at 100 kb resolution on hg38. Compartment switches with FDR ≤ 0.05 were considered significant. To enable integration with our T2T-CHM13–based gene and RTE annotations, hg38 compartment calls were lifted over to T2T-CHM13 coordinates using (Hinrichs et al. 2006). For each gene, the compartment status of its overlapping bin was used to classify it as residing in a stable A, stable B, switching toward A, or switching toward B region. Concordance between compartment switching and differential expression was assessed by intersecting compartment-switching genes with differentially expressed genes (FDR ≤ 0.05); concordant genes were those whose direction of expression change matched the predicted direction of compartment switching (toward A and upregulated, or toward B and downregulated). Over-representation analyses were performed using fgsea (Korotkevich et al. 2016).

Over-representation analyses of curated senescence-associated gene sets (Hallmark Interferon Alpha Response, SenMayo, Reactome Extracellular Matrix Organization, KEGG Lysosome, GOBP Response to ER Stress, Reactome Cell Cycle) used Fisher’s exact test on the intersection of each gene set with each compartment-switching category. Venn diagrams were drawn using eulerr (Larsson 2021).

### Subcompartment Analysis

Subcompartments were called at 25 kb resolution on hg38 using Calder 2 (the hg38-compatible release of Calder) (Liu et al. 2021), which assigns each bin to one of eight subcompartments ranging from A.1.1 (most active) to B.2.2 (most repressed). Subcompartment transitions were defined as changes in subcompartment assignment between conditions, with the number of “jumps” calculated as the signed difference along the A.1.1 → B.2.2 axis. As with compartments, subcompartment calls were lifted over from hg38 to T2T-CHM13 coordinates (Hinrichs et al. 2006) for integration with downstream gene and RTE annotations. Genes overlapping subcompartment-switching regions were identified using bedtools intersect (Quinlan and Hall 2010) with default settings. Concordance with differential expression and gene ontology analyses were performed as described in the Compartment Analysis section.

### Loop Calling and Cross-Condition Comparison

Loops were called from each merged Hi-C map using Juicer’s CPU implementation of HiCCUPS (Durand et al. 2016) with default parameters. To compare loop sets across conditions, we developed a custom matching script. For each pair of conditions, two loops were considered the same if both anchors of one loop fell within ±n bp of the corresponding anchors of the other loop (Figure S4A). The threshold n was selected by sweeping values from 10 kb to 100 kb on chromosome 21 and identifying the value at which the count of unique and shared loops plateaued; n = 30 kb was used for all genome-wide comparisons (Figure S4B,C). Loop graph visualizations were generated with NetworkX (Hagberg et al., n.d.). Loop anchor compartment status, CTCF site overlap, and methylation status were calculated for each anchor. Genes were classified by their loop status across conditions (shared, gained, or lost) and intersected with differential expression results.

### RNA-seq Library Preparation and Sequencing

Total RNA was extracted from proliferating and RS LF1 cells using TRIzol. Quiescent cell pellets were submitted directly. All samples were sent to Azenta for library preparation using a stranded ribosomal-depletion protocol. Proliferating and RS libraries were sequenced across two HiSeq lanes, and quiescent libraries on one HiSeq lane.

### RNA-seq Core Analysis

We analyzed these data with our TE-Seq pipeline (Kelsey et al. 2025) and custom scripts. Full TE-Seq pipeline methods can be found at https://doi.org/10.1186/s13100-025-00381-w. Briefly, sequencing reads were trimmed with Fastp (Chen et al. 2018) and aligned to a sample-specific TE-patched genome (see Methods: Custom LF1 Reference Genome Generation) using STAR (Dobin et al. 2013). Gene expression was quantified using featureCounts (Liao et al. 2014) and a RefSeq transcript annotation (GCF_009914755.1). Repetitive element expression was quantified using the telescope tool (Bendall et al. 2019) in both “unique” and “multi” modes, which consider either uniquely mapped reads or both uniquely and multi-mapping reads. Genes were assessed for differential expression using DESeq2 (Love et al. 2014). We used GSEA to query MSigDB gene sets for enrichment (Subramanian et al. 2005). Differential expression analysis of aggregated RTE subfamily counts was performed using a negative binomial generalized linear model (GLM). For this analysis, raw counts were summed across group members, and sample size factors were incorporated as an offset in the model. For locus-resolved RTE analyses, loci were intersected with chromatin architectural features (compartment status, subcompartment switching, loop membership) using the GenomicRanges R package (Lawrence et al. 2013). Genome-occupancy versus expression-share comparisons (Figure 3B) were performed by computing, for each compartment class, the fraction of family-level genomic copies and the fraction of family-level expression originating from that compartment.

### DNA Extraction, Library Preparation, and Sequencing

We isolated genomic DNA using the NEB Monarch Spin gDNA Extraction Kit. Sequencing libraries were prepared following the unmodified Nanopore DNA Ligation Sequencing Kit V14 (SQK-LSK114) protocol for a PromethION flow cell. Each sample was loaded onto its own 10.4.1 PromethION Flow Cell. Sequencing was performed on a P2-solo instrument, which allows two flow cells to be run concurrently. Two sequencing libraries were prepared for each sample. Midway through sequencing, flow cells were washed with the flow cell wash kit (EXP-WSH004-XL) and reloaded with a fresh library from the same sample. Sequencing was controlled with MinKNOW software version 25.03.9 and run for three days or until nearly all sequencing pores were inactive. Four proliferating and four RS samples were sequenced, yielding ∼28x mean coverage and ∼20 kb N50 per sample.

### Nanopore Basecalling and Quality Control

Nanopore sequencing data were basecalled using Dorado (VN:0.9.5+bd7fb217) with the super-accuracy, 5mCG and 5hmCG basecalling model (dna_r10.4.1_e8.2_400bps_sup@v5.0.0_5mCG_5hmCG@v3). Basecalled reads were then aligned to the T2T-CHM13 (GCF_009914755.1) human genome (Nurk et al. 2022). Quality control metrics were generated using PycoQC (Leger and Leonardi 2019). We then created a filtered BAM for sensitive downstream applications requiring only high-confidence alignments. Using samtools (Li et al. 2009), we retained only alignments with a read quality score ≥ 10 and a MAPQ ≥ 10 and removed secondary and supplementary alignments (samtools view –b –F 2304 –q 10 –e ‘[qs] >= 10’). Loop anchor methylation differences (Figure 2D) were calculated as the mean per-anchor methylation difference (RS – PRO), with anchors stratified by class (proliferation-unique, shared, RS-unique). CTCF motif methylation profiles (Figure 2E) were generated by extracting per-CpG methylation values within ±1 kb of all CTCF motif centers (JASPAR PWM scan; see below) overlapping loop anchors of each class, then averaging across motifs and replicates. For locus-level methylation at L1HS_14q23.2_3 (Figure 5), per-CpG methylation across the locus and ±10 kb of flanking sequence was computed and smoothed with a rolling mean. Per-CpG methylation values across the L1 5′UTR were extracted in 5′→3′ order. Single-read methylation maps were generated with WGBS tools (Loyfer et al. 2026) by visualizing the per-CpG methylation status of individual Nanopore reads spanning the full L1HS_14q23.2_3 5′UTR.

### Custom LF1 Reference Genome Generation

We used our TE-Seq pipeline (Kelsey et al. 2025) to call non-reference RTE insertions and create sample-specific, non-reference TE-patched genomes. Full methods can be found at https://doi.org/10.1186/s13100-025-00381-w. Briefly, we aligned Nanopore DNA sequencing reads from LF1 genomic DNA to the T2T-CHM13 genome using Minimap2 (Li 2018). Approximately 230x coverage of pooled Nanopore DNA reads was provided as input to TLDR (Ewing et al. 2020), which identifies and characterizes private insertions. Insertion “patches” comprising each private insert sequence plus 30 bp of flanking sequence on either side of the breakpoint were appended to the T2T-CHM13 reference (Nurk et al. 2022) as additional contigs. We annotated these contigs for repetitive element content with RepeatMasker (“RepeatMasker Home Page,” n.d.) and merged them with the T2T-CHM13 reference RepeatMasker annotations. All reference and private L1HS and L1PA2 elements were inspected for intact protein-coding potential by extracting their sequences with bedtools getfasta (Quinlan and Hall 2010), detecting open reading frames with the ORFik R package (Tjeldnes et al. 2021), and filtering on length to identify elements retaining intact ORF1p and ORF2p coding capacity. These elements were classified as “intact.”

### CTCF Motif and Binding Site Analysis

Sequence-based CTCF motif positions were defined by scanning the hg38 genome for matches to the JASPAR CTCF position weight matrix (Rauluseviciute et al. 2023) using FIMO from the MEME Suite (Bailey et al. 2009) with default parameters, and lifted over to T2T-CHM13 coordinates (Hinrichs et al. 2006). To complement these motif-based predictions with empirical binding data, we incorporated ENCODE (ENCODE Project Consortium 2012) CTCF ChIP-seq peak calls from IMR-90 primary lung fibroblasts (file ENCFF203SRF, biosample ENCBS379ENC), lifted over from hg38 to T2T-CHM13 (chm13v2.0) in the same way. Loop anchors overlapping either a ChIP-seq peak or a motif site were considered CTCF-associated. For methylation profiles around CTCF sites, ChIP-seq peaks were centered on their summits, and motif sites not overlapping a peak were added, centered on their midpoints.

### RT-qPCR Validation in IMR-90 Cells

For RT-qPCR validation experiments (Figure S6B), IMR-90 female human diploid lung fibroblast cells were used. IMR-90 cells were maintained in DMEM with 10% FBS and 1% Penicillin/Streptomycin/Glutamine, at 37°C, 5% CO₂, and 2.5% O₂. Proliferating cells were harvested at 60–80% confluency. Contact-inhibited quiescent cells were obtained by allowing cells to reach 100% confluency and maintaining them at confluency for 4 days in regular growth media. Serum-withdrawal quiescent cells were obtained by switching cells at 60% confluency to growth media containing 0.01% FBS for 4 days. Etoposide-induced senescent cells were generated by treating cells with 40 µM etoposide in regular growth media every other day for two weeks. RNA was extracted using the QIAGEN RNeasy micro kit and reverse transcribed using the QIAGEN QuantiTect Reverse Transcription Kit. qPCR was performed on the resulting cDNA using Applied Biosystems Power SYBR Green PCR Master Mix on an Applied Biosystems QuantStudio 6 Pro Real-Time PCR System according to the manufacturer’s instructions. Primer sequences for senescence and SASP genes (CDKN1A, CDKN2A, IL1B, IL6, IL8, IFNA) were as previously published (De Cecco et al. 2019). PUM1 was used as the reference gene, as previously validated (González-Bermúdez et al. 2019).

## Declarations

### Ethics approval and consent to participate

Not applicable

### Consent for publication

Not applicable

### Availability of data and materials

Hi-C and RNA-seq data (both raw and processed – including compartment, subcompartment, and loop calls) can be found GEO accession numbers GSE268486 and GSE268487. Nanopore DNA-sequencing data have been deposited in the Sequencing Read Archive (SRA), BioProject accession number PRJNA1375768. Code can be found at: https://github.com/maxfieldk/Senescence_HiC

The most up-to-date version of the TE-Seq pipeline (which was used for sample-specific, non-reference TE-patched genome generation and RNA-seq analysis) can be found at: https://github.com/maxfieldk/TE-Seq, where instructions for pipeline installation, configuration, and deployment are documented.

Any other data or information relevant to this study are available from the corresponding author upon reasonable request.

### Competing interests

JMS is a cofounder and SAB chair of Transposon Therapeutics, cofounder of GeroSen Biotechnologies, and serves as consultant to RoC Skincare. All other authors declare they have no competing interests.

### Funding

This work was supported in part by the following NIH grants: R01AG050582 (NN, JMS), UG3CA268202/UH3CA268202 (NN), UM1HG011593 (NN), RM1HG011016 (NN), R01AG016694 (JMS), R01AG078925 (JMS), P01AG051449 (JMS, MMGK), T32AG041688 (AD, AR, TAN), T32GM128596 (AD), T32GM007601 (AD, AR), F31AG072748 (AR). It was also supported by a Blavatnik Family Fellowship to MMGK and to AD.

### Author contributions

Conceptualization: MMGK, AD, SAE, JMS, NN, Methodology: AD, MMGK, SAE, TAN, AR, KC, JMS, NN, Investigation: MMGK, AD, SAE, TAN, AR, KC, Visualization: MMGK, AD, Supervision: JMS, NN, Writing – original draft: MMGK, AD, NN, Writing – review & editing: MMGK, AD, SAE, TAN, AR, KC, JMS, NN

## Acknowledgements

The authors are thankful to all members of the Neretti Lab, Sedivy Lab, Dr. Erica Larschan, and members of the CEGS Center for Genome Imaging for fruitful discussion. We are grateful to the Brown Center for Computation and Visualization (CCV) team for their management of the OSCAR high performance computing cluster which was used throughout this work.

---

**Figure S1.**
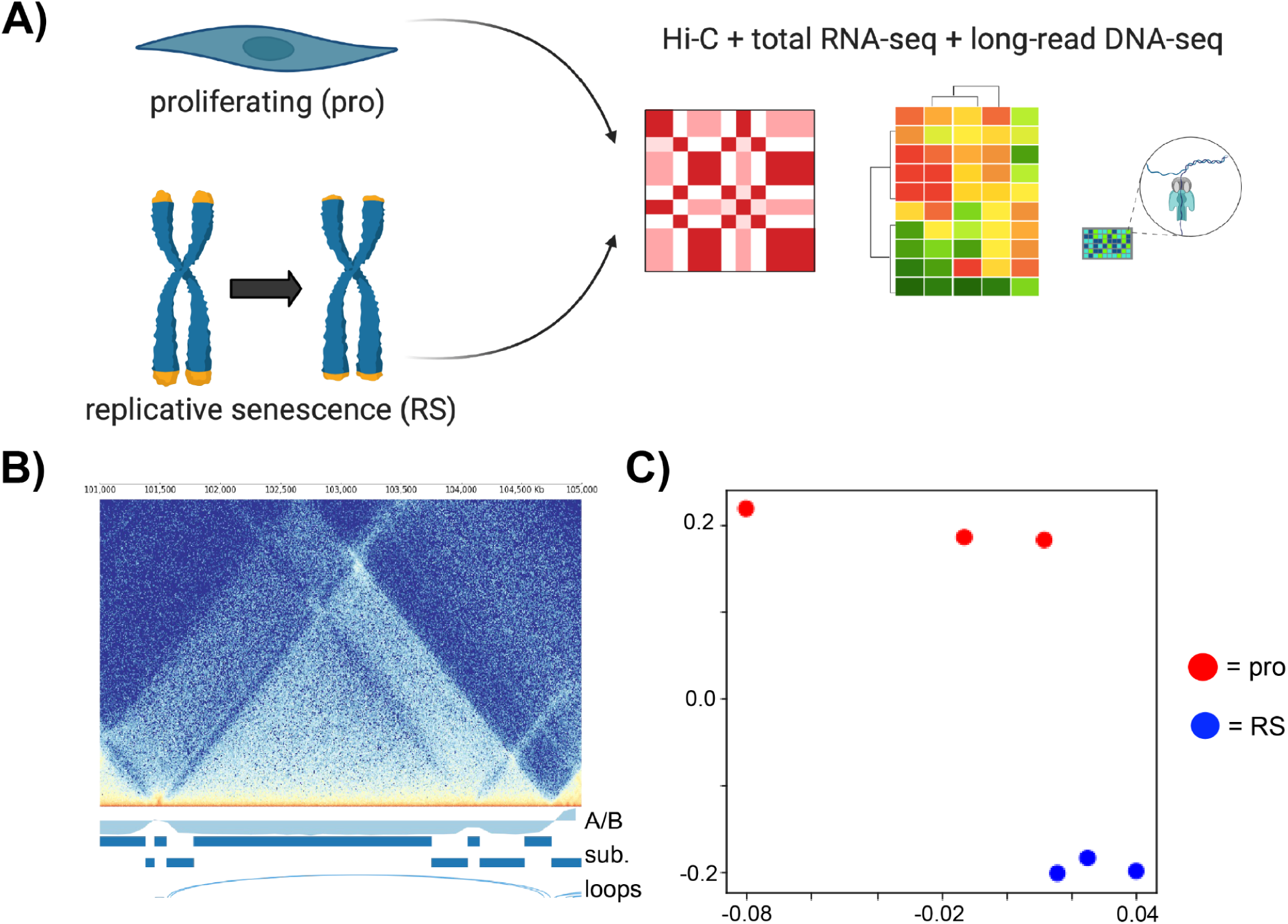
| Quality control and experimental overview for Hi-C maps. (**A**) Schematic overview of the experimental workflow. Hi-C, total RNA-seq, and long-read DNA sequencing were performed on proliferating and late-stage replicatively senescent (RS) LF1 fibroblasts. (**B**) Representative high-resolution Hi-C contact map displaying key architectural features and genomic annotations. (**C**) Multidimensional scaling (MDS) plot showing tight clustering of biological replicates for both proliferating and RS conditions, indicating high reproducibility of Hi-C datasets.

**Figure S2.**
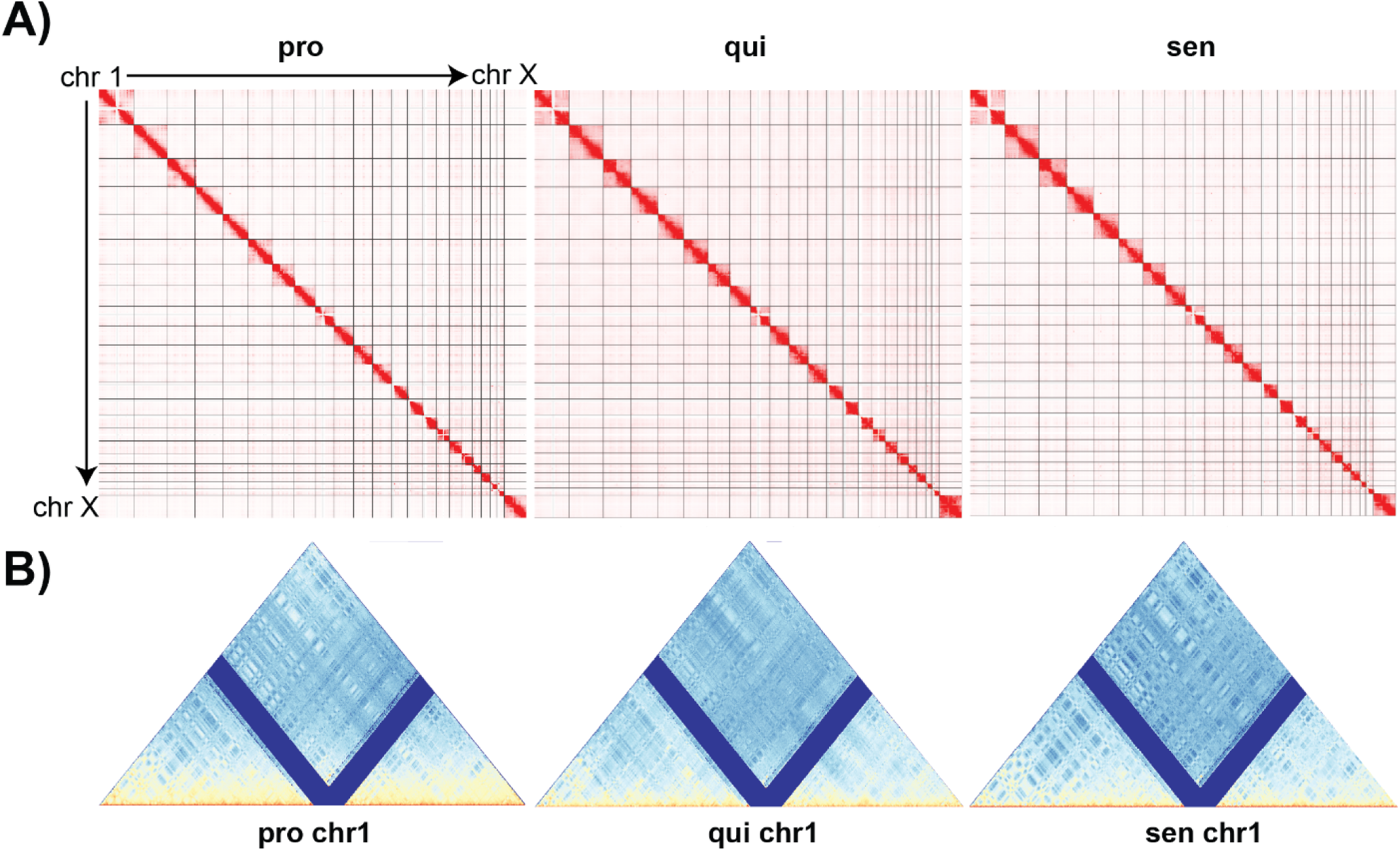
| Full genome Hi-C visualizations. (**A**) Full-genome Hi-C maps of proliferating, quiescent, and senescent cells. (**B**) Chromosome 1 Hi-C map for proliferating, quiescent, and senescent cells.

**Figure S3.**
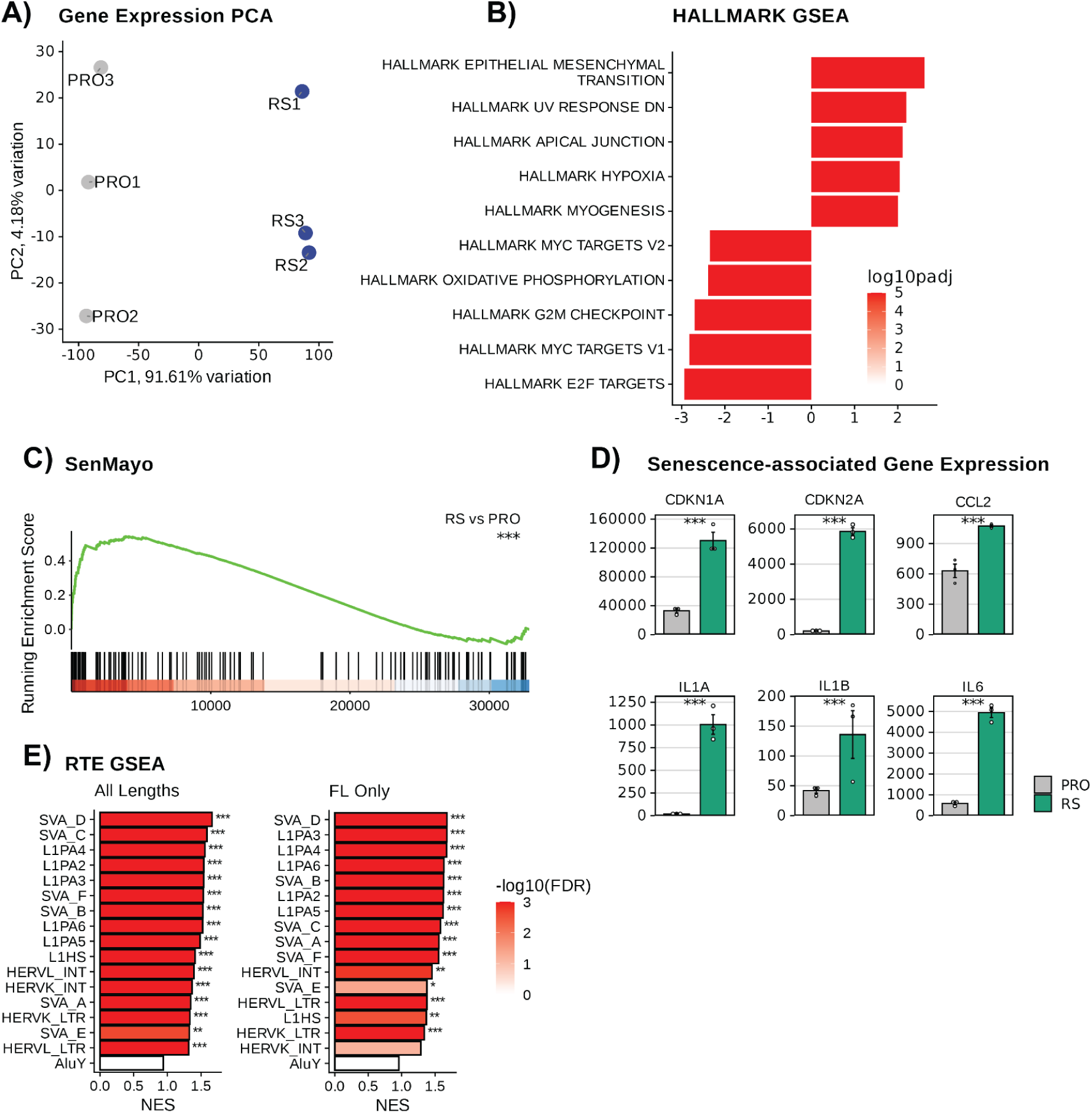
| Total RNA-seq confirms canonical senescence transcriptional signatures in RS LF1 fibroblasts. (**A**) Principal component analysis of total RNA-seq from proliferating (PRO) and replicatively senescent (RS) LF1 fibroblasts. PC1 (91.61% variation) cleanly separates RS from PRO. (**B**) Hallmark GSEA of RS versus PRO (ranked by DESeq2 Wald statistic). Bars show normalized enrichment scores (NES); color reflects –log10 adjusted p-value. (**C**) GSEA running enrichment plot for the SenMayo senescence signature (RS vs. PRO; FDR < 0.001). (**D**) Expression of senescence-associated and SASP transcripts. Bars show mean ± SE; *** denotes FDR < 0.001 (DESeq2). (E) Pre-ranked GSEA of LINE-1 subfamily enrichment in senescence (RS vs. PRO). All RepeatMasker-annotated elements served as the ranked background (DESeq2 Wald statistic). Left: all elements per subfamily; right: full-length (FL) elements only. Bars show normalized enrichment score (NES); fill indicates −log10(FDR); *, FDR < 0.05; **, FDR < 0.01; ***, FDR < 0.001 (fgsea).

**Figure S4.**
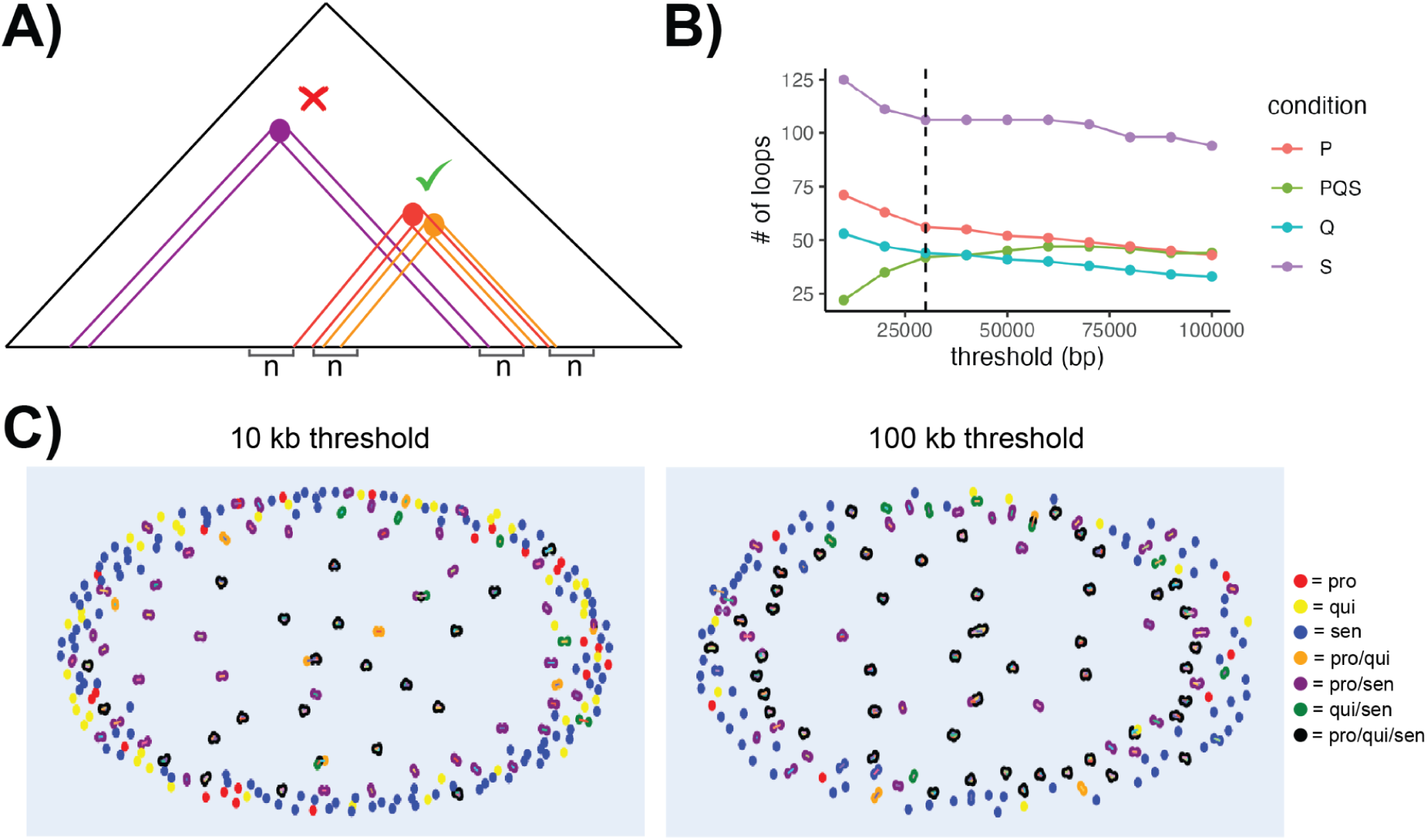
| Strategy and parameter selection for cross-condition loop matching. (**A**) Schematic of the loop-matching procedure. Two loops from different conditions are classified as the same (green check) if both anchors fall within ±n base pairs of one another; otherwise they are classified as condition-unique (red ×). (**B**) Threshold sweep on chromosome 21 across n values from 10 kb to 100 kb, showing the number of loops assigned to each class — proliferating-unique (P), quiescent-unique (Q), RS-unique (S), and shared across all three conditions (PQS) — as a function of n. The dashed vertical line marks n = 30 kb, the threshold adopted for genome-wide analyses, where loop-class counts plateau. (**C**) Loop graph visualizations of chromosome 21 loops at the 10 kb (left) and 100 kb (right) thresholds, illustrating how the choice of n affects the partitioning of loops into condition-unique and shared classes. Nodes represent loops; node color indicates condition membership (red = pro-only, yellow = qui-only, blue = sen-only, orange = pro/qui, purple = pro/sen, green = qui/sen, black = pro/qui/sen).

**Figure S5.**
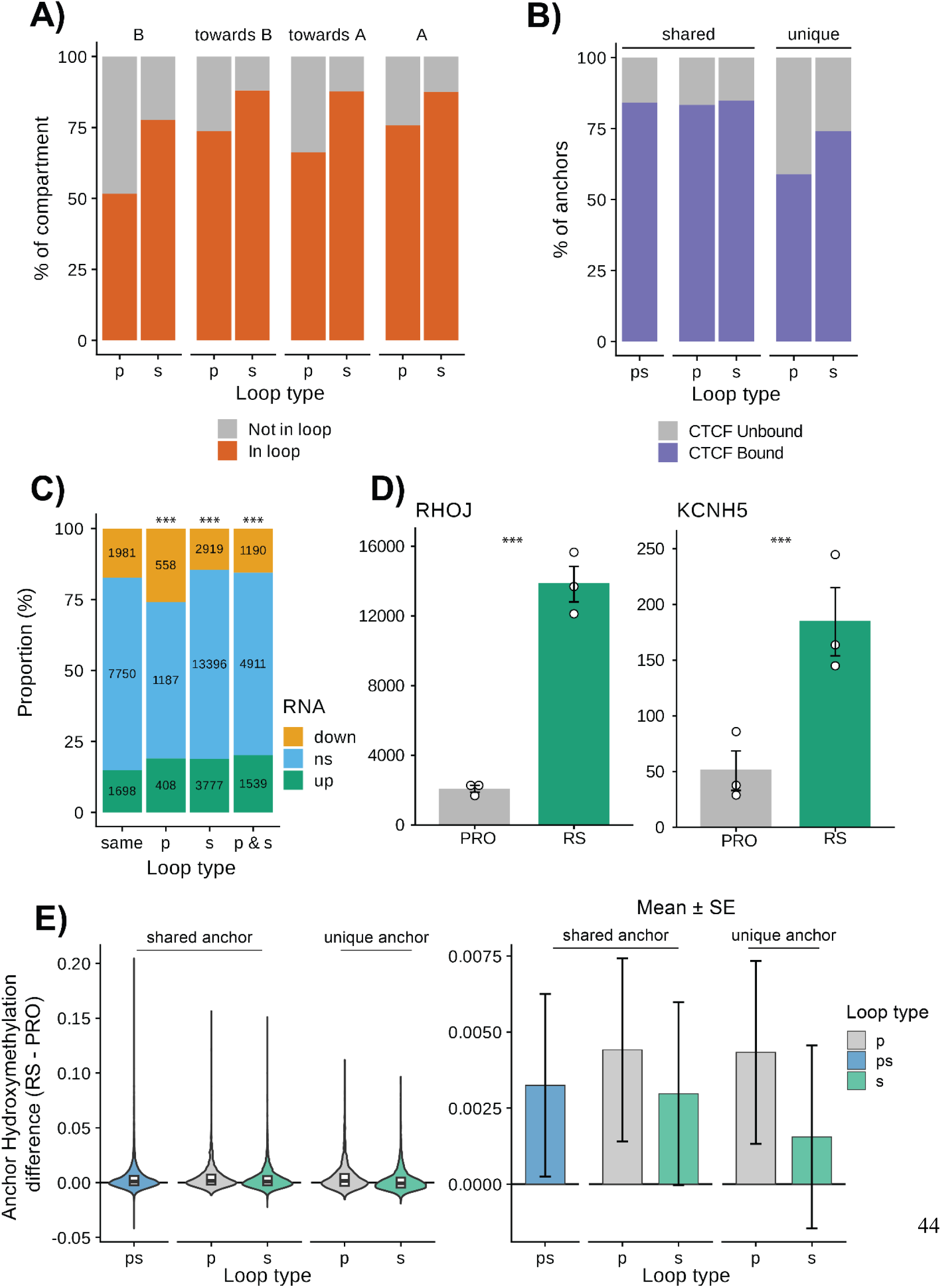
| Loop occupancy of compartments, CTCF anchor coverage, and concordance between loop status and differential expression. (**A**) Percentage of each compartment class engaged in any chromatin loop in proliferating (p) and RS (s) cells, stratified by stable B, switching toward B, switching toward A, and stable A regions. (**B**) Percentage of loop anchors overlapping a CTCF site (ENCODE IMR-90 CTCF ChIP-seq peak or CTCF motif), stratified by anchor class. Each loop contributes its two anchors. Unique anchors overlapped CTCF sites less often than shared anchors (Fisher’s exact test, BH-adjusted p < 0.001 for all comparisons). (**C**) Differential expression status (RS vs. PRO) of genes stratified by their loop status – whether this is stable across conditions (same), or whether they have lost one or more proliferation-unique loops in senescence (p), have gained one or more rs-unique loops in senescence (s), or have both lost and gained at least one loop in senescence (p & s). Numbers indicate gene counts per category. Genes which gain or lose loop contacts between conditions are more likely to be differentially expressed than genes with stable loop status (Fisher’s exact test, p ≤ 0.001 for all comparisons). (**D**) Expression of L1HS_14q23.2_3 s-loop associated genes (RHOJ, KCNH5). Bars show mean ± SE; *** denotes BH-adjusted p < 0.001 (DESeq2). (**E**) CpG Hydroxymethylation difference (RS – PRO) at loop anchors, by anchor class. Left: per-anchor distributions. Right: linear model-estimated methylation difference (RS − PRO) ± SE.

**Figure S6.**
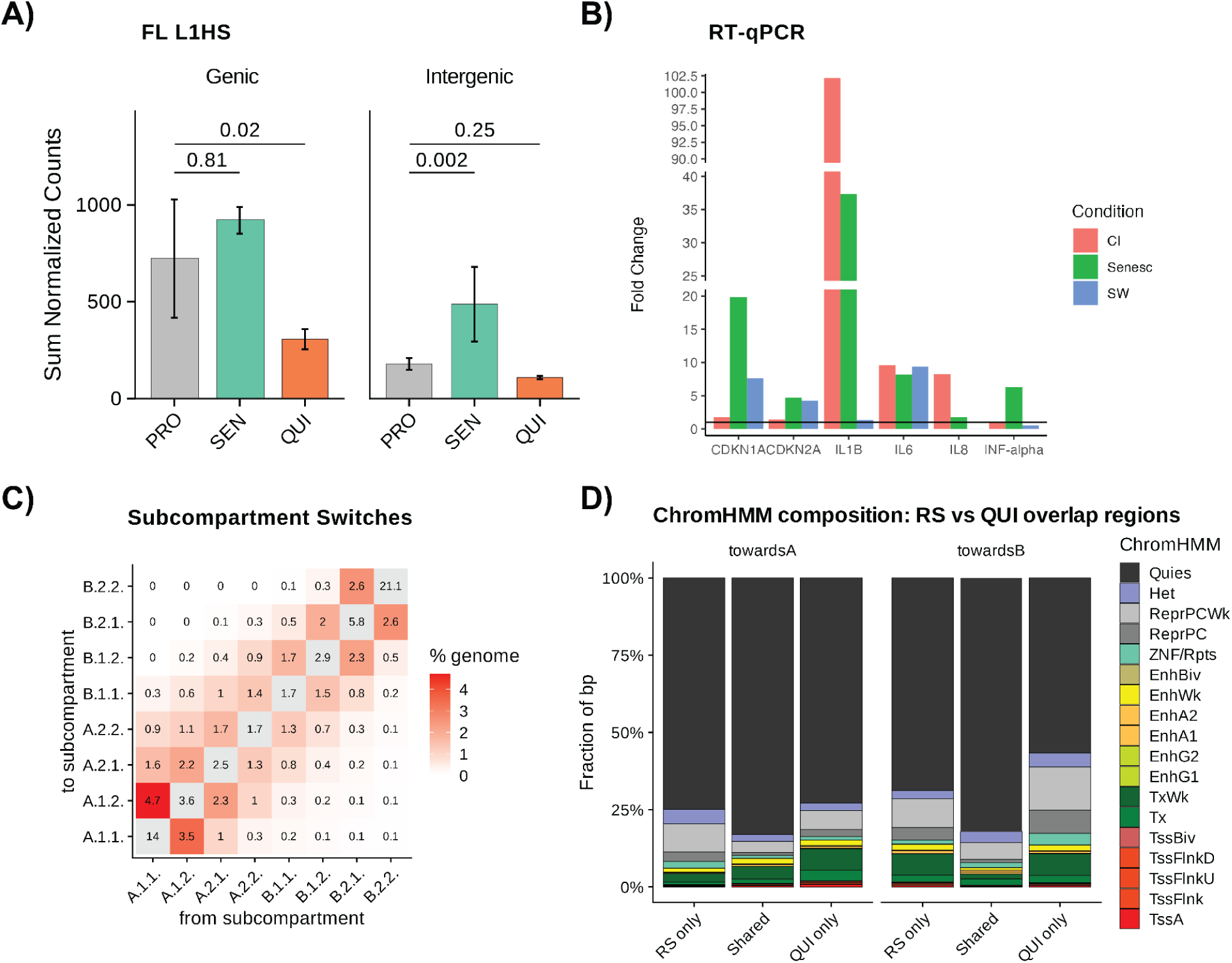
| Quiescence shows distinct subcompartment dynamics from RS but shares a senescence-like inflammatory transcriptional response across induction protocols. (A) Sum of normalized expression counts across FL L1HS elements, stratified by genic context, in proliferating (PRO), replicatively senescent (RS), and quiescent (QUI) cells. Bars show mean ± SE. Statistical significance was assessed using a negative binomial GLM. (B) RT-qPCR assessment of senescence and SASP genes in IMR-90 fibroblasts under contact inhibition (CI), serum withdrawal (SW), and etoposide-induced senescence (Senesc). Fold change is relative to proliferating controls; PUM1 served as the reference gene. (C) Percentage of the genome undergoing each subcompartment transition from PRO (columns) to QUI (rows). (D) ChromHMM state composition of RS-only, shared, and QUI-only regions undergoing compartment switching toward A (left) or toward B (right). Color indicates ChromHMM state; bar height shows fraction of base pairs.

**Table S1.** Number of sequenced reads and resulting Hi-C contacts per replicate.

| **Sample Name** | **Replicate** | **Sequenced Reads** | **Hi-C Contacts** |
|---|---|---|---|
| Proliferating | 1 | 479,606,053 | 335,144,966 |
| Proliferating | 2 | 376,935,533 | 263,471,076 |
| Proliferating | 3 | 394,567,767 | 274,118,940 |
| Quiescent | 1 | 453,732,287 | 286,795,901 |
| Quiescent | 2 | 444,787,457 | 281,594,189 |
| Quiescent | 3 | 479,962,618 | 300,708,434 |
| Senescent | 1 | 508,313,932 | 316,312,022 |
| Senescent | 2 | 466,055,552 | 294,431,607 |
| Senescent | 3 | 449,124,662 | 304,263,231 |

## Notes

### Competing Interest Statement

JMS is a cofounder and SAB chair of Transposon Therapeutics Inc., holds equity in PrimeFour and Atropos Therapeutics, and consults for Atropos Therapeutics and Longaevus Technologies. All other authors declare they have no competing interests.

### Summary of Updates

The manuscript has been substantially revised to incorporate new experimental data, additional computational analyses, and clarifications of the findings and their interpretation. We generated new high coverage Nanopore long read sequencing data from proliferating and replicatively senescent LF1 fibroblasts, with four biological replicates per condition. These data confirm preferential DNA hypomethylation at senescence specific chromatin loop anchors, while showing that CTCF binding motifs remain protected. Additional analyses of DNA hydroxymethylation and CTCF binding further clarify the relationship between DNA methylation and chromatin loop formation. We expanded the analyses of chromatin compartment switching, gene expression, and retrotransposon activation, including new gene set enrichment analyses. We also added higher resolution chromatin interaction maps and quantitative loop measurements for a specific LINE1 element, together with detailed analysis of its promoter methylation.

